# Large-scale geography survey provides insights into the colonization history of a major aphid pest on its cultivated apple host in Europe, North America and North Africa

**DOI:** 10.1101/2020.12.11.421644

**Authors:** S.G. Olvera-Vazquez, C. Remoué, A. Venon, A. Rousselet, O. Grandcolas, M. Azrine, L. Momont, M. Galan, L. Benoit, G. M. David, A. Alhmedi, T. Beliën, G. Alins, P. Franck, A. Haddioui, S.K. Jacobsen, R. Andreev, S. Simon, L. Sigsgaard, E. Guibert, L. Tournant, F. Gazel, K. Mody, Y. Khachtib, A. Roman, T.M. Ursu, I.A. Zakharov, H. Belcram, M. Harry, M. Roth, J.C. Simon, S. Oram, J.M. Ricard, A. Agnello, E. H. Beers, J. Engelman, I. Balti, A. Salhi-Hannachi, H. Zhang, H. Tu, C. Mottet, B. Barrès, A. Degrave, J. Razmjou, T. Giraud, M. Falque, E. Dapena, M. Miñarro, L. Jardillier, P. Deschamps, E. Jousselin, A. Cornille

## Abstract

With frequent host shifts involving the colonization of new hosts across large geographical ranges, crop pests are good models for examining the mechanisms of rapid colonization. The microbial partners of pest insects may also be involved in or affected by colonization processes, which has been little studied so far. We investigated the demographic history of the rosy apple aphid, *Dysaphis plantaginea*, a major pest of the cultivated apple (*Malus domestica*) in Europe, North Africa and North America, as well as the diversity of its microbiota. We genotyped a comprehensive sample of 714 colonies from Europe, Morocco and the US using mitochondrial (*CytB* and *CO1*), bacterial (16s rRNA and *TrnpB*), and 30 microsatellite markers. We detected five populations spread across the US, Morocco, Western and Eastern Europe and Spain. Populations showed weak genetic differentiation and high genetic diversity, except the ones from Morocco and North America that are likely the result of recent colonization events. Coalescent-based inferences revealed high levels of gene flow among populations during the colonization but did not allow determining the sequence of colonization of Europe, North America and Morroco by *D. plantaginea*, likely because of the weak genetic differentiation and the occurrence of gene flow among populations. We found that *D. plantaginea* rarely hosts other endosymbiotic bacteria than its obligate nutritional symbiont *Buchnera aphidicola*. This suggests that secondary endosymbionts did not play an important role in the rapid spread of the rosy apple aphid. These findings have fundamental importance for understanding pest colonization processes and implications for sustainable pest control programs.

## Introduction

Understanding the evolutionary processes underlying the colonization of new environments and the range expansion of species is a key goal in evolutionary biology (Angert, Bontrager, & Ågren, 2020; Austerlitz, Jung-Muller, Godelle, & Gouyon, 1997; Excoffier, Foll, & Petit, 2009; Hoffmann & Courchamp, 2016; Rius & Darling, 2014). Crop pests in agro-ecosystems, with their frequent colonization of new hosts across large geographic ranges, are good models to study the mechanisms of rapid colonization and range expansion (Garnas et al., 2016; Gladieux et al., 2014; Stukenbrock & McDonald, 2008). The key questions relating to the evolutionary processes underlying the colonization, spread and success of crop pests pertain to the geographic origin of the source population, the location of the colonization routes, the extent to which genetic diversity is reduced via founder effects (Blakeslee et al., 2019) and the extent of gene flow among populations during the spread of the plant parasite (Stukenbrock, 2016). Current and future threats to biodiversity and their consequences on ecosystem health and services make these questions more relevant than ever. Understanding the routes of pest colonization contributes greatly to the efforts to protect crops against future pest emergence and therefore has direct implications for breeding and agronomic programs that develop biological methods of parasite control (Estoup & Guillemaud, 2010; Fraimout et al., 2017; Lawson Handley et al., 2011; Turcotte, Araki, Karp, Poveda, & Whitehead, 2017).

Historically, gene genealogies have been a rich source of information into a species’ evolutionary history that could be applied to the study of colonization (Bloomquist, Lemey, & Suchard, 2010; Hickerson et al., 2010; Posada & Crandall, 2001). The characterization of population structure, genetic diversity and demographic history (divergence time, migration rates among populations and effective population size) are also essential to understand the evolutionary processes underlying rapid colonization and range of geographic expansion (Excoffier et al., 2009). Approximate Bayesian computation methods (referred to as “ABC” hereafter) provide a robust framework for inferring a species’ history by allowing the comparison of alternative demographic models and the estimation of their associated parameters (divergence time, migration rate, effective population size) (Bertorelle, Benazzo, & Mona, 2010; Csilléry, Blum, & Francois, 2012; Csilléry, Blum, Gaggiotti, & François, 2010; Estoup, Raynal, Verdu, & Marin, 2018; Raynal et al., 2019; Roux & Pannell, 2015). The power of the ABC methods has made it possible to retrace the evolutionary history of notorious plant parasites (*e*.*g*., *Plasmopara viticola* (Berk & Curtis) Berl. & de Toni (Fontaine et al., 2020), *Microbotryum lychnidis-dioicae* (DC. Ex Liro) G. Deml & Oberw. (Gladieux et al., 2015)), insect crop pests (*e*.*g*., *Batrocera dorsalis* Hendel (Aketarawong et al., 2014), *Drosophila suzukii* Matsumura (Fraimout et al., 2017), *Daktulosphaira vitifoliae* Fitch (Rispe et al., 2020)) and invasive alien species posing a threat to the native fauna (*e*.*g*., *Harmonia axyridis* Pallas (Lawson Handley et al., 2011)). These studies have identified the source populations, reconstructed complex colonization routes, and determined the pace of geographic range expansion, often emphasizing the role of human transportation in the spread of these noxious species. Recently, the ABC approach, combined with machine learning (*i*.*e*., random forest, referred to as “ABC-RF” hereafter (Estoup et al., 2018; Raynal et al., 2019)), was used to demonstrate that the African arid-adapted locust pest species *Schistocerca gregaria* Forsskål colonized Africa through major migration events driven by the last glacial climatic episodes (Chapuis et al., 2020). Yet, ABC methods are underused for estimating the extent of gene flow during parasite colonization (but see (Fraimout et al., 2017)). Most studies assume punctual admixture events among populations, but rarely continuous gene flow among populations. Only recently has the new ABC-RF approach been used to infer the invasion routes, evolutionary history and extent of gene flow in the spotted-wing *D. suzukii* Matsumura from microsatellite markers (Fraimout et al., 2017) (referred to as SSR for simple sequence repeat hereafter). Beyond population genomics approaches, and in the special case of insect pests, the investigation of colonization history could also benefit from the characterization of insect endosymbiotic bacterial communities. Indeed, many insect pests host a consortium of endosymbiotic bacteria that mediate their adaptation to new environmental conditions (Frago, Zytynska, & Fatouros, 2020). Variations in the endosymbiotic consortium along colonization routes could facilitate the rapid adaptation of insect pests to different environments (Lenhart & White, 2020) and, in consequence, its spread. Alternatively, if the colonization stems from only a few populations, it might be accompanied by a loss of endosymbiont diversity along the colonization routes.

Aphids are a good study system to investigate the evolutionary processes involved in range expansion and the colonization of new environments. Aphids infest a wide range of host species and can be major pests of many crop plants (Blackman & Eastop, 2000). Some aphid species have now become cosmopolitan following the dissemination of crops around the globe (Brady et al., 2014; Kirk, Dorn, & Mazzi, 2013; Zepeda-Paulo et al., 2010; Zhang, Edwards, Kang, & Fuller, 2014). The clonal reproduction of aphids during spring and summer is one of the reasons put forward to account for their remarkable success worldwide. Indeed, asexual reproduction allows for a rapid increase in population size after the colonization of a favorable new environment (Simon, Rispe, & Sunnucks, 2002; Simon, Stoeckel, & Tagu, 2010; Figueroa, Fuentes-Contreras, Molina-Montenegro, & Ramírez, 2018). So far, only a handful of studies have reconstructed the colonization history of aphid crop pests by combining population genetics approaches using the information from SSR, sequence or single nucleotide polymorphism (SNP) markers (Fang, Chen, Jiang, Qu, & Qiao, 2018; Giordano et al., 2020; Kim, Hoelmer, & Lee, 2016; Leclair, Buchard, Mahéo, Simon, & Outreman, 2021; Morales-Hojas, Sun, Iraizoz, Tan, & Chen, 2020; Peccoud et al., 2008; Piffaretti et al., 2013; Zepeda-Paulo et al., 2010; Zhang et al., 2014, 2014; Zhou et al., 2015). These studies demonstrated that aphid species can spread very quickly across the world, probably via plants transported by humans and/or wind. These investigations detected a colonization involving several populations with high genetic diversity, possibly with gene flow (Wang, Hereward, & Zhang, 2016; Wei, Zuorui, Zhihong, & Lingwang, 2005) and/or a few “super-clones”, *i*.*e*., predominant genotypes widespread in space and time (Vorburger, Lancaster, & Sunnucks, 2003; Piffaretti et al., 2013; Figueroa, Fuentes-Contreras, Molina-Montenegro, & Ramírez, 2018). Assessing the genetic diversity, genetic structure and the extent of gene flow among populations in aphids are therefore central to determining the evolutionary processes that have occurred during aphid colonization.

Associating the reconstruction of aphid colonization history with the characterization of their endosymbiotic bacterial community can shed light on the processes of their dispersal. Aphids harbor both obligate symbionts that supply them with the nutrients missing from their diet (Buchner, 1965) and facultative symbionts that can provide various selective advantages in specific environmental conditions (Haynes et al., 2003; Oliver, Degnan, Burke, & Moran, 2010). The obligate aphid endosymbiont bacterium *Buchnera aphidicola* has strictly codiverged with most aphid species (Jousselin, Desdevises, & Coeur d’acier, 2009). Bacterial markers (*e*.*g*., *TrpB*), along with other typical markers (*e*.*g*., *CO1* or *Cytb*), can be used to infer the phylogenetic history of aphid species and to investigate signals of recent range expansions (Popkin et al., 2017; Zhang et al., 2014). Beyond the use of bacterial genomes to help reconstruct aphid phylogeography, the composition of the bacterial populations in aphids might also help to assess the importance of facultative bacteria for colonizing new geographic regions. Many studies have investigated variation in bacterial communities associated with the pea aphid (*Acyrthosiphon pisum* Harris), revealing geographical variation, host-specific differentiation and associations with environmental factors such as temperature, host plants, and natural enemies (Russell et al., 2013; Tsuchida, Koga, Shibao, Matsumoto, & Fukatsu, 2002; Zepeda-Paulo, Ortiz-Martínez, Silva, & Lavandero, 2018; Leclair et al., 2021). The effect of endosymbionts on aphid fitness has been confirmed experimentally in certain cases (Frago et al., 2017; Leclair et al., 2016). Nevertheless, studies on a global scale in non-model aphid species are still scarce (Zytynska & Weisser, 2016), and there are as of yet no studies simultaneously investigating the colonization routes of an aphid species and the changes in symbiotic associations along this route.

*Dysaphis plantaginea* Passerini, the rosy apple aphid, is one of the most harmful aphid pests attacking cultivated apple trees (*Malus domestica* Borkh), causing major economic losses every year, especially in Europe, North Africa and North America (Guillemaud, Blin, Simon, Morel, & Franck, 2011; Warneys et al., 2018; Wilkaniec, 1993). This aphid species occurs across temperate regions (Central and Southwest Asia, North Africa, North America, and Europe) (Blackman & Eastop, 2000) in cultivated apple orchards. The rosy apple aphid completes its life cycle on two successive host plants: the cultivated apple trees as its sole primary host plant, from early autumn to late spring, and the plantain herb *Plantago* spp. as a secondary host plant during summer (Bonnemaison, 1959). The rosy apple aphid reproduces through cyclical parthenogenesis whereby clonal reproduction alternates with a sexual reproduction, the latter taking place in autumn when females lay fertilized overwintering eggs on apple trees. Eggs hatch in early spring (Blommers, Helsen, & Vaal, 2004). While its phylogenetic relationships with other aphid species are quite well resolved, the evolutionary history of *D. plantaginea* has been little explored to date (but see (Guillemaud et al., 2011)). The native geographical range of *D. plantaginea* and its ancestral host range are not known. It might have been associated with *Malus sieversii*, the primary ancestor of the cultivated apple (*M. domestica*) (Harris, Robinson, & Juniper, 2002), and then colonized Europe during the journey of the cultivated apple along the Silk Routes from Asia to Europe. Alternatively, the rosy apple aphid may have colonized its cultivated apple host in Europe rapidly and recently, about 1,500 years ago when the Greeks brought the cultivated apple to Europe from Central Asia (Cornille et al., 2019; Cornille, Giraud, Smulders, Roldán-Ruiz, & Gladieux, 2014). These scenarios are derived from our knowledge of the domestication history of apples; there are no data on the rosy apple aphid that would support any of these scenarios. The colonization routes of the rosy apple aphid are unknown, except those historical records document the introduction of the rosy apple aphid in North America was recent (*ca*. 1890s) (Foottit, Halbert, Miller, Maw, & Russell, 2006). More generally, the population structure and the extent of gene flow among populations throughout the geographic distribution of *D. plantaginea* are largely unknown, including the regions where it causes the most damage in apple orchards, *i*.*e*., North America, North Africa and Europe. There are also no data so far regarding the diversity of endosymbionts across a large geographical range in aphids. Here, we investigate the colonization history of this major fruit tree pest using multiple approaches and genetic datasets, drawn from comprehensive samples taken from the primary host, the cultivated apple in Europe, North America and Morocco. Note that, despite repeated attempts, we failed to collect *D. plantaginea* in its putative source region where apple trees originated in Central Asia, preventing us from fully addressing its earliest colonization history. Therefore, we aimed to answer the following questions focusing on regions most negatively affected by the rosy apple aphid, *i*.*e*., Europe, North America and North Africa: i) What is the spatial genetic diversity and population structure of *D. plantaginea* across Europe, the US and Morocco? Can we detect genetically differentiated populations, and/or recent bottlenecks in colonizing populations? ii) Did gene flow occur among populations during colonization? iii) Did *D. plantaginea* populations lose or gain symbionts during their colonization?

## Methods

### Samples and DNA extraction

Each sample described hereafter consisted of a single aphid colony of 10-15 females collected on a host plant during the spring of 2017 and 2018. Sampling only one colony per tree ensured that the sample did not contain different clones, which can occur on the same tree. Each colony was kept in ethanol (96%) at −20°C until DNA extraction. For the three methods described below (*i*.*e*., SSR genotyping, Sanger sequencing of the aphid mitochondrial *CO1, Cytb* markers, and *TrpB* bacterial marker, and metabarcoding of the 16S rRNA bacterial marker), DNA was isolated from a single individual per colony using a new standardized protocol (Supplementary material Text S1). We used different individuals to obtain DNA for the amplification of the different markers. For the sake of simplicity, colonies are referred to as ‘individual’ or ‘samples ‘hereafter. Note that samples used for 16S rRNA sequencing underwent two extra chemical washes before DNA extraction to remove the external bacteria that could be present on the aphid’s cuticle. The extra chemical washes consisted of a first wash with dithiothreitol/DTT (50 mM) for 4 minutes, followed by a second wash with potassium hydroxide/KOH (200 mM) for 4 minutes. The KOH wash was performed twice. Five negative controls were also included ((Jousselin et al., 2016), Table S1).

Different sample sizes were used for each of the three methods (*i*.*e*., SSR genotyping, metabarcoding of the 16S rRNA bacterial marker, Sanger sequencing of the aphid *CO1, Cytb* and *TrpB* markers). For each sample, the locality, sample collector identity, host plant species, latitude, longitude and use in this study (genotyping, Sanger sequencing and/or metabarcoding) are given in Table S1.

The largest sample was collected for SSR genotyping, comprising 667 *D. plantaginea* samples (colonies) from Europe, Morocco and the US, from three hosts: *M. domestica* (50 sites, *i*.*e*., orchards, *N* = 654), but also *M. sylvestris* (one site, Alta Ribagorça in Spain, *N* = 7), the European wild apple, and *P. lanceolata*, the secondary host (one site, Loos-en-Gohelle in France, *N*=6). The 667 samples originated from 52 different geographic sites (*i*.*e*., 52 orchards) spread over 13 countries; seven to 15 individuals were collected at each site (Table S1, Figure S1). We tried to obtain samples from Eastern Asia and Central Asia during fieldwork in 2017 and 2018, and through our collaborative network. However, despite our attempts and although *D. plantaginea* is referenced in the literature on various hosts in several Central Asian countries (Aslan & Karaca, 2005; CAB International, 2020; Holman, 2009), it was not observed in these areas.

For the investigation of the bacterial 16S rRNA region, we used 178 *D. plantaginea* individuals out of the same 667 colonies used for SSR genotyping (Table S1). We selected two to three samples (colonies) per site to cover a wide and even spatial distribution across Europe and North America. The selected 178 individuals were collected across 12 countries on *M. domestica*, except eight on *M. sylvestris* (Table S1). Morocco was the only country not represented for the bacterial 16S rRNA analysis.

For Sanger sequencing of *CO1, TrpB* and *CytB*, we used a total of 84 samples belonging to eight aphid species (*D. plantaginea, Dysaphis* sp., *Aphis citricola* van der Goot, *Aphis pomi* de Geer, *Aphis spiraecola* Patch, *Melanaphis pyraria* Passerini, *Myzus persicae* Sulzer, *Rhopalosiphum insertum* Walker) sampled on five host plant species (*M. domestica, M. sylvestris, Sorbus aucuparia, Prunus persica, Pyrus communis*). The species were chosen to represent the aphid genera known to be closely related to the rosy apple aphid (Choi, Shin, Jung, Clarke, & Lee, 2018). One to three individuals from each of the geographic sites listed above and in Table S1 were sampled, for a total of 67 samples for *D. plantaginea*. The 17 additional samples from seven other aphid species (one to two samples per aphid species) were collected on the cultivated apple and other fruit tree species (Table S1).

### PCR and Sanger sequencing

We amplified the coding regions from the aphid mitochondrial cytochrome c oxidase subunit I (*CO1*) gene and the cytochrome B (*CytB*) gene, as well as the Tryptophan synthase subunit B (*TrpB*) from *Buchnera*. Fragments were amplified following the protocol reported in Popkin et *al*. (2017) with some modifications (Table S2). The final PCR volume was 30 μL, containing Buffer (1X), MgCl_2_ (1.5 mM), dNTP (0.1 mM), Forward and Reverse primers (0.7 μM), 5 μL of Taq polymerase (1 U), and 5 μL of DNA (1/10 dilution). Amplification products were visualized on an agarose gel 1.5% stained with ethidium bromide under ultraviolet light. We prepared four 96-well plates with 15 μL of PCR products and a negative control. Plates were sent to Eurofins Genomics France SAS for sequencing.

Chromatograms were inspected and corrected manually with CodonCode Aligner version 8.0.1 (www.codoncode.com), assigned as an ‘N’ when two peaks overlapped. Alignment, evaluation of all coding genes for frameshifts and elimination of pseudogenes were performed with MEGA version 7.0.26 (Kumar, Stecher, & Tamura, 2016). The neutral evolution of each gene, and thus its suitability for phylogenetic and population genetic analyses, was assessed with the McDonald and Kreitman test (Egea, Casillas, & Barbadilla, 2008; McDonald & Kreitman, 1991) (Table S3). Two samples of *Brachycaudus helichrysi* Kaltenbach, a pest of *Prunus*, for which sequences of *CO1* (NCBI sequence identifiers: KX381827.1, KX381828.1), *CytB* (KX381989.1, KX381990.1), and *TrpB* (KX382153.1, KX38215.1) were available (Popkin et al., 2017), were added in the phylogenetic analyses. Sequences were concatenated, resulting in a data matrix of 86 concatenated sequences.

### Phylogenetic tree and taxonomic assignation

We checked the taxonomic assignation of the samples used in this study by running phylogenetic analyses including *D. plantaginea* and other aphid species found on fruit trees with a Bayesian approach implemented in MrBayes v3.2.7 (Huelsenbeck & Ronquist, 2001) and Randomized Axelerated Maximum Likelihood-RAxML (Stamatakis, 2014), using the GTR “Generalized time-reversible” mutational model. The two *M. pyraria* individuals were used as an external group. We chose the default parameters (unlink statefreq = (all) revmat = (all) shape = (all); prset applyto = (all) ratepr = variable; mcmcp ngen = 1000000 nruns = 2 nchains = 4 samplefreq = 1000 printfreq = 1000). Inferred trees were visualized with FigTree v1.4 (http://tree.bio.ed.ac.uk/software/figtree/).

### SSR genotyping

We used 30 SSR markers, including one that was previously used for *D. plantaginea* (Guillemaud et al., 2011), and 29 that were newly developed from the sequencing of a low coverage genome (see details of the protocol in supplementary material Texts S2 and S3). We tested the neutrality, and the absence of linkage disequilibrium, of the 29 SSR markers using the Ewens-Watterson neutrality test ((Watterson, 1978); Text S3 and Table S4). Each SSR was amplified separately by PCR. PCR was performed in a final volume of 20 µL (0.2 µM of each forward and reverse primer, with the forward primer labeled with a fluorescent dye, 0.2 µM of dNTPs, between 1 and 1.5 mM of MgCl_2_, 1X Buffer (5X), 2 µL of a homemade Taq, 5 µL of DNA (1/30 dilution) and sterile H_2_O to reach the final volume). We used the following PCR program: 94°C for 5 minutes, followed by 35 cycles of 30s at 94°C, 30s at 60 to 65°C and 45s at 72°C, then 5 minutes at 72°C and finally 10 minutes at 4°C. In the PCR program, the annealing temperature varied between 55°C and 65°C. Annealing temperatures for each SSR marker are detailed in Table S5. PCR products were then pooled according to the four multiplexes described in Table S5.

SSR genotyping was performed at the GENTYANE platform (INRAE, Clermont-Ferrand, France). Alleles of each SSR marker were identified, and their size scored with Genemapper v.4.0 (Applied Biosystems TM, Foster City, USA) by two people independently. In case of discrepancy, the electropherogram was triple checked for a final decision. Allele scoring resulting from Genemapper® was then processed with the Autobin Excel Macro (https://www6.bordeaux-aquitaine.inra.fr/biogeco_eng/Scientific-Production/Computer-software/Autobin). We retained only multilocus genotypes with less than 30% missing data and containing less than 5% null alleles. Null alleles were detected with GENEPOP v4.7 (Rousset, 2008).

### Clonal population structure

Each individual was classified according to its multilocus genotype (*MLG*) with GenoDive 2.0b23 ((Meirmans & Van Tienderen, 2004), Table S1). We used the stepwise mutation model with a threshold of 0 and the corrected Nei’s diversity index as a statistic to test clonal population structure. To reduce the influence of clonal copies produced by asexual reproduction on Hardy–Weinberg equilibrium, allele frequency and genetic differentiation estimates, the dataset was pruned to include only one copy of each *MLG* for further analyses.

### Population genetics descriptive statistics

Observed and expected heterozygosity (*H*_*O*_ and *H*_*E*_) and inbreeding coefficient (*F*_*IS*_) were calculated with GENEPOP v4.7 (Rousset, 2008) from the SSR dataset for each SSR marker, each site (*i*.*e*., geographic location/orchard) and each population (*i*.*e*., clusters inferred with the STRUCTURE software, to comprise individuals with a membership coefficient of at least 62.5% to the given cluster, see results). The 62.5% cut-off was chosen based on the distribution of individual membership coefficients across the clusters detected for the most likely *K* value (see results). Pairwise genetic differentiation (*F*_*ST*_) between sites and between populations was also calculated with GENEPOP (Rousset, 2008). Only sites with at least five successfully genotyped samples were included for the site-specific computations. Allelic richness and private allelic richness for each site and each population were calculated with ADZE (Szpiech, Jakobsson, & Rosenberg, 2008) using a sample size corresponding to the smallest number of observations per site or population, multiplied by two chromosomes (*e*.*g*., a sample size of 20 represents ten individuals x two chromosomes).

We also estimated Nei’s nucleotide diversity index *π* (Nei, 1987), Watterson’s index q (Watterson, 1975), haplotype diversity, Tajima’s *D* (Tajima, 1989) and Fu’s *Fs* (Fu, 1997) with DNAsp (Rozas, Sànchez-DelBarrio, Messeguer, & Rozas, 2003) using the concatenated three-marker dataset (*i*.*e*., *CO1, TrpB* and *CytB*) for each population (*i*.*e*., cluster inferred with STRUCTURE including individuals with membership coefficient > 62.5% to this cluster, see results).

### Detecting recent bottlenecks during population range expansion

We tested whether a bottleneck occurred during the range expansion of each population with the method implemented in BOTTLENECK (Cornuet & Luikart, 1996; Piry, Luikart, & Cornuet, 1999). Inferences regarding historical changes in population size are based on the principle that the expected heterozygosity estimated from allele frequencies decreases faster than the expected heterozygosity estimated under a given mutation model at mutation-drift equilibrium in populations that have experienced a recent reduction in size. The tests were performed under the stepwise-mutation model (SMM) and a two-phase model (TPM) allowing for 30% multi-step changes.

### Spatial distribution of allelic richness and observed heterozygosity

Spatial patterns of allelic richness and observed heterozygosity were visualized by mapping the variation in allelic richness and observed heterozygosity at 48 sites in total (*i*.*e*., sites with at least five individuals) with the geometry-based inverse distance weighted interpolation in QGIS (Quantum GIS, GRASS, SAGA GIS). The correlation between genetic variability (*H*_*O*_) and latitude, and between *H*_*O*_ and longitude, was tested using a linear model.

### Population subdivision

We inferred the finest population structure by comparing the results obtained with three population genetic tools: STRUCTURE v2.3.2 (Pritchard, Stephens, & Donnelly, 2000), TESS v2.3.1 (Chen, Durand, Forbes, & François, 2007), and a discriminant analysis of principal components (DAPC) (Jombart, Devillard, & Balloux, 2010). STRUCTURE is based on the use of Markov chain Monte Carlo (MCMC) simulations to infer the assignment of genotypes to *K* distinct clusters. In addition to this, TESS also considers a spatial component, so that genotypes from sites that are geographically closer to each other are considered more likely to be in the same cluster. For both STRUCTURE (using admixture model with correlated allele frequencies) and TESS (using hierarchical mixture model), ten independent analyses were carried out for each value of *K* (1 ≤ *K* ≤ 10 and 2 ≤ *K* ≤ 10, respectively) with 500,000 MCMC iterations after a burn-in of 50,000 steps. STRUCTURE and TESS outputs were processed with CLUMPP 1.1.2 (Jakobsson & Rosenberg, 2007) to identify potential distinct modes (*i*.*e*., clustering solutions) in replicated runs (10) for each *K*. We also assessed the population subdivision with DAPC with the R package ‘adegenet’ (Jombart & Collins, 2015), which does not rely on any assumption about the underlying population genetics model, in particular concerning Hardy-Weinberg equilibrium or linkage equilibrium. The number of genetic clusters was investigated with the *find*.*cluster* function (Jombart & Collins, 2015; Ripley & Ripley, 2001), which runs successive *K*-means for clustering. The automatic cluster selection procedure ‘*diffNgroup*’ was used with *n*.*iter* set to 10^6^ and *n*.*start* set to 10^3^. The ordination analysis (DAPC) was performed using the *dapc* function. The statistically optimal number of principal components was assessed using the *optim*.*a*.*score* function. Assessment of the samples assigned to a genetic cluster was performed using the *compoplot* function.

We used Pophelper (Francis, 2016) to run the Evanno method on the STRUCTURE outputs. The Evanno method detects the strongest level of population subdivision (Evanno, Regnaut, & Goudet, 2005). For TESS, we used the rate of change of the deviation index criterion (*DIC*) to determine the amount of additional information explained by increasing *K*. For DAPC, we looked at the Bayesian information criterion (*BIC)* obtained with the *adegenet* package to estimate the optimal *K* value. However, the *K* identified with the *DIC, BIC* and *ΔK* statistics does often not correspond to the finest biologically relevant population structure (Cornille et al., 2015; Kalinowski, 2011; Puechmaille, 2016). We therefore visualized the bar plots with Pophelper (Francis, 2016) and chose the *K* value for which all clusters had well assigned individuals while no further well-delimited and biogeographically relevant clusters could be identified for higher *K* values. For further analyses, we considered an individual to be assigned to a cluster when its membership coefficient was ≥ 62.5% to this cluster (see results below).

The spatial pattern of genetic structure was visualized by mapping the mean membership coefficients for each site, as inferred from each of the three population genetics structure analyses, with QGIS 3.12 ‘Las Palmas ‘(https://qgis.org). We further explored relationships among populations with a principal component analysis performed on a table of standardized alleles frequencies (PCA, dudi.pca, ade4 R package (Dray & Dufour, 2007)).

### Isolation-by-distance

We tested whether there was a significant isolation-by-distance (IBD) pattern. A Mantel test with 10,000 random permutations was performed between the individual coefficient of relatedness *F*_*ij*_ (Loiselle, Sork, Nason, & Graham, 1995) and the matrix of the natural logarithm of geographic distance. We also performed a correlation between *F*_*ST*_/(1-*F*_*ST*_) and the natural logarithm of geographic distance. These analyses were performed using SPAGeDI 1.3 (Hardy & Vekemans, 2002) separately for each *D. plantaginea* panmictic population (*i*.*e*., cluster containing individuals with a membership coefficient > 0.625) identified with TESS, STRUCTURE, and DAPC.

### Demographic and divergence history using ABC-RF

We used ABC to investigate whether the spatial patterns of genetic clustering, diversity and differentiation observed in *D. plantaginea* resulted from the occurrence of gene flow among populations during colonization. We also attempted to infer the sequence of colonization events in each population. We used the recently developed ABC method based on a machine learning tool named “random forest” (ABC-RF) to perform model selection and parameter estimations (Estoup et al., 2018; Pudlo et al., 2016; Raynal et al., 2019). In brief, this method creates a “forest” of bootstrapped decision trees to classify scenarios based on the summary statistics of the datasets. Some simulations are not used to build the trees and can thus be used to cross-validate the analysis by computing a prior error rate. This approach allows the comparison of complex demographic models (Pudlo et al., 2016) by comparing groups of scenarios with a specific type of evolutionary event with other groups with different types of evolutionary events (instead of considering all scenarios separately) (Estoup et al., 2018).

We used a nested ABC approach with two key steps. First, we inferred the divergence and demographic history of the rosy apple aphid in Europe (step 1). Then, we tested the divergence and demographic history of the rosy apple aphid outside of Europe (step 2). Each ABC step compared different sequences of colonization events, with and without bidirectional gene flow among populations (Figure S2). This two-step nested approach avoids the need to compare models that are too complex, which would require the simulation of too many populations and parameters and is more powerful than testing all scenarios individually to determine the main evolutionary events that characterize demographic history and divergence (Estoup et al., 2018). Populations were defined as the clusters detected with STRUCTURE (see results), removing putative admixed individuals (*i*.*e*., individuals with a membership coefficient < 0.625 to any given cluster). The model parameters used were the divergence time between *X* and *Y* populations (*T*_*X-Y*_), the effective population size of population *X* (*N*_*E-X*_), the migration rate per generation between *X* and *Y* populations (*m*_*X-Y*_). Prior values for divergence time were drawn for the log-uniform distribution bounded between the distributions used in the ABC and are given in Table S6.

For all models, identical SSR datasets were simulated for 29 out of the 30 markers that had perfect repeats (we excluded the L4 marker because it did not have perfect repeats, Tables S4 and S5), increasing confidence in the simulated model. We preliminarily checked that the population structure inferred with 29 SSR markers did not differ significantly from the inferences obtained with 30 SSR markers (data not shown). We assumed a generalized stepwise model of SSR evolution. Mutation rates were allowed to vary across loci, with locus-specific mutation rates drawn from a gamma distribution (*α, α/μ*) where *μ* is the mutation rate per generation and α is a shape parameter. We assumed a log-uniform prior distribution for *μ* (1e-4, 1e-3) and a uniform distribution for *α* (1.30).

We used ABCtoolbox (Wegmann, Leuenberger, Neuenschwander, & Excoffier, 2010) with fastsimcoal 2.5 (Excoffier & Foll, 2011) to simulate datasets, using model parameters drawn from prior distributions (Table S6). We performed 10,000 simulations per scenario how has been suggested (Pudlo et al., 2016). For each simulation, we calculated six summary statistics per population with arlsumstats v3.5 (Excoffier and Lischer 2010): *H*, the mean heterozygosity across loci, *sd(H)*, the standard deviation of the heterozygosity across loci, *GW*, the mean Garza-Williamson statistic across loci (Garza and Williamson, 2001), *sd(GW)*, the standard deviation of the mean Garza-Williamson statistic over populations, *NGW*, the mean modified Garza-Williamson statistic over loci, *sd(NGW)*, the standard deviation of the mean modified Garza-Williamson statistic over populations. We also computed pairwise *F*_*ST*_ (Weir and Cockerham, 1984) and genetic distances (*δμ*)^2^ (Goldstein et al., 1995) between pairs of populations.

We used the abcrf v.1.7.0 R statistical package (Pudlo et al., 2016) to carry out the ABC-RF analysis. This analysis provides a classification vote that represents the number of times a scenario is selected as the best one among *n* trees in the constructed random forest. For each ABC step, we selected the scenario, or the group of scenarios, with the highest number of classification votes as the best scenario, or best group of scenarios, among a total of 500 classification trees (Breiman, 2001). We then computed the posterior probabilities and prior error rates (*i*.*e*., the probability of choosing a wrong group of scenarios when drawing model index and parameter values from the priors of the best scenario) over 10 replicate analyses (Estoup et al., 2018). We also checked visually that the simulated models were compatible with the observed dataset by projecting the simulated and the observed datasets onto the two first linear discriminant analysis (LDA) axes (Pudlo et al., 2016), and checking that the observed dataset fell within the clouds of simulated datasets.

We calculated parameter inferences using the final selected model following the two-step ABC procedure. Note that the ABC-RF approach includes the model checking step that was performed *a posteriori* in previous ABC methods.

### Characterization of the bacterial community associated with the rosy apple aphid using the 16S rRNA bacterial gene

In order to investigate the bacterial diversity in *D. plantaginea* populations, we amplified a 251 bp portion of the V4 region of the 16S rRNA gene (Mizrahi-Man, Davenport, & Gilad, 2013) and used targeted sequencing of indexed bacterial fragments on a MiSeq (Illumina) platform (Kozich, Westcott, Baxter, Highlander, & Schloss, 2013) following the protocol described in (Jousselin et al., 2016). We used 178 aphid DNA extracts (Table S1), comprising 175 *D. plantaginea* individuals and three *M. pyraria* individuals (Table S1). We wanted to represent a range as large as possible from our samples. We also added eight randomly chosen samples that did not undergo the two extra chemical washes (see Materials and methods, and Table S1). Each sample was amplified twice along with negative controls (DNA extraction and PCR controls). PCR replicates were conducted on distinct 96-well microplates. We obtained a total of 390 PCR products (186 DNA extracts, by 2 for PCR duplicates, plus PCR controls), which were pooled and then separated by gel electrophoresis. Bands based on the expected size of the PCR products were excised from the gel, purified with a PCR clean-up and gel extraction kit (Macherey-Nagel), and quantified with the Kapa Library Quantification Kit (Kapa Biosystems). Paired-end sequencing of the DNA pool was carried out on a MISEQ (Illumina) FLOWCELL with a 500-cycle Reagent Kit v2 (Illumina).

We first applied sequence filtering criteria following Illumina’s quality control procedure. We then used a pre-processing script from (Sow et al., 2019) to merge paired sequences into contigs with FLASH V.1.2.11 (Magoč & Salzberg, 2011) and trim primers with CUTADAPT v.1.9.1 (Martin, 2011). We then used the FROGS pipeline (Escudié et al., 2018) to generate an abundance table of bacterial lineages across samples. In brief, we first filtered out sequences > 261 bp and < 241 bp with FROGS, then we clustered variants into operational taxonomic units (OTUs) with SWARM (Mahé, Rognes, Quince, Vargas, & Dunthorn, 2014) using a maximum aggregation distance of three. We identified and removed chimeric variants with VSEARCH (Rognes, Flouri, Nichols, Quince, & Mahé, 2016). We only kept OTUs that were present in both PCR replicates of the same sample and then merged the number of reads for each OTU for each aphid sample.

Taxonomic assignment of OTUs was carried out using RDPtools and Blastn (Altschul, Gish, Miller, Myers, & Lipman, 1990) against the Silva138-16s database (https://www.arb-silva.de) as implemented in FROGS. From the abundance table of OTUs across samples, we transformed read numbers per aphid sample into frequencies (percentages); sequences accounting for < 0.5 % of all the reads for a given sample were excluded following Jousselin e*t al*. (2016). All filters resulted in excluding reads found in low abundance that could represent sequencing errors and which were also often found in the negative controls.

## Results

### Taxonomic status of aphid samples

The Bayesian phylogenetic tree built with the 86 sequences representing nine aphid species resulted in a polytomy for the 67 *D. plantaginea* samples from Europe, Morocco and the USA (Figure S3), showing very little sequence variation at the intraspecific level (low bootstrap values < 0.6, Figure S3a). We therefore kept only two representatives out of the 67 *D. plantaginea* individuals in subsequent phylogenetic analyses. Bayesian analysis of this pruned dataset comprising 19 individuals, representing nine aphid species, confirmed known phylogenetic relationships in Aphidinae ((Choi et al., 2018), Figure S3b). The species *A. pomi, A. citricola* and *A. spiraecola* were grouped in the same clade, the three species appearing polyphyletic (Figure S3b). *Dysaphis plantaginea* appeared closely related to other representatives of the Macrosiphini tribe (*B. helichrysi* and *M. persicae*). Interestingly, the samples from Iran clustered apart from the *D. plantaginea* clade, the former belonging to the *Dysaphis* sp. species. This result suggests that the two *Dysaphis* sp. Iranian samples belong to a yet unidentified species or population exhibiting strong differentiation from European populations. Those two samples were not included in the population genetics analyses using SSR, as we only had two representatives from Iran (Table S1).

### Clone detection

Overall, the proportion of unique genotypes was variable among sites. We found 582 unique genotypes, among which 29 were repeated (12.7% of the total dataset, Table S7) and involved 85 individuals, mainly coming from Belgium (86% of the clones, eight sites; mean proportion of unique genotypes (mean G/N) = 0.36 ± 0.2), Bulgaria (6%, one site, G/N = 0.67), France (5%, three sites, mean G/N = 0.87 ± 0.05), the USA (1 %, one site; G/N = 0.93), Spain (1%, one site; G/N = 0.91) and from the lab-reared aphids (1%, G/N = 0.09). We kept only the 582 unique genotypes for the analyses presented below.

### Spatial distribution of allelic variation

The map of allelic richness (Figure 1a) showed that genetic diversity decreased along a northeast to southwest gradient, with the highest allelic richness found in northeastern Europe and the lowest in Morocco and the USA, except for Belgium that showed a lower level of genetic diversity. We found a significant correlation between allelic richness and longitude (*r* = 0.229, *P-value* = 0.001), as well as between allelic richness and latitude (*r* = 0.268, *P-value* = 0.001). The map of observed heterozygosity (Figure 1b) confirmed that genetic diversity decreased along a northeast to southwest gradient, with the highest allelic richness in Denmark and the lowest in Morocco. We found a significant correlation between observed heterozygosity and longitude (*r* = 0.173, *P-value* = 0.003), and between observed heterozygosity and latitude (*r* = 0.499, *P-value* = 1.95e-08).

**Figure 1.**
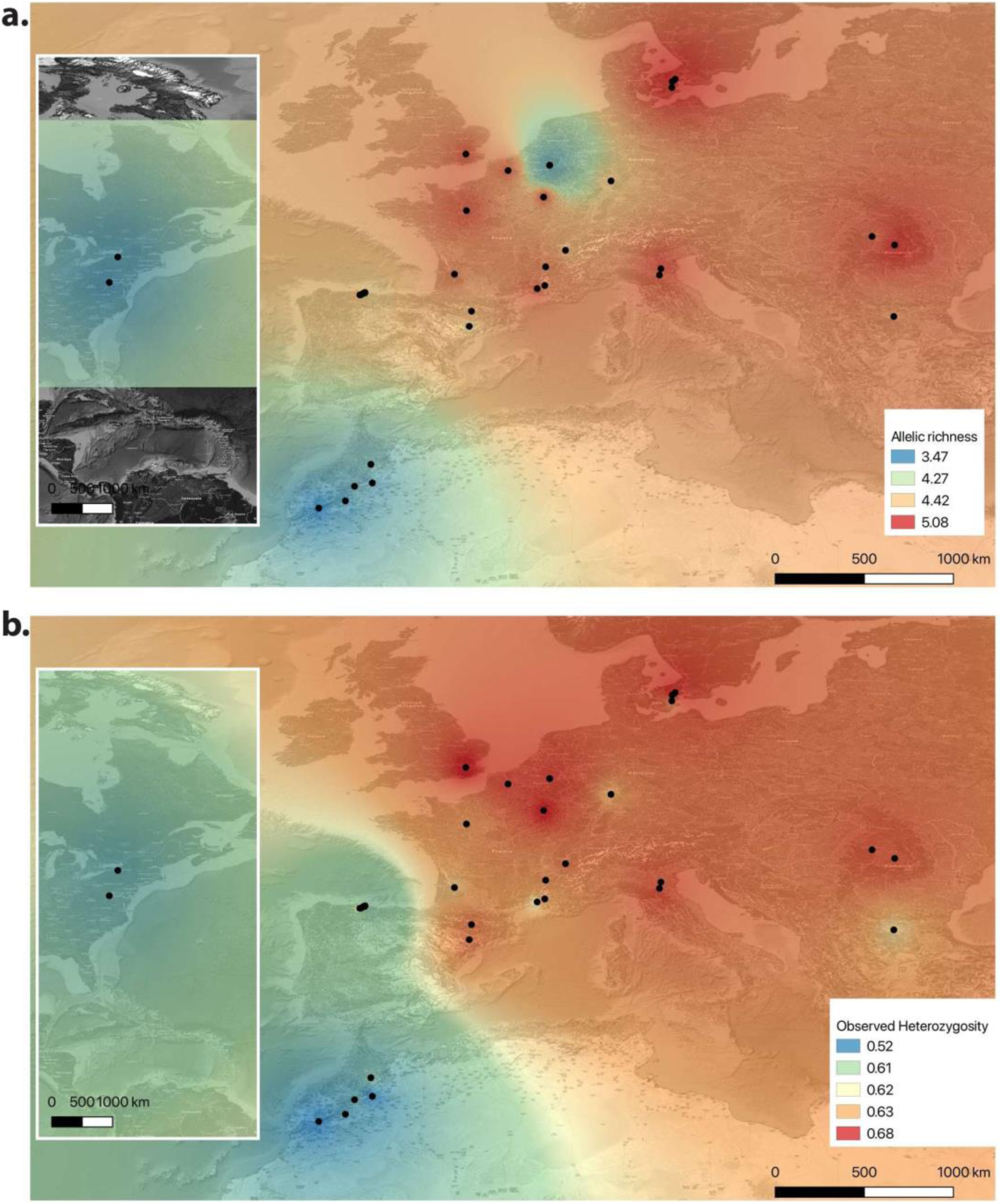
Spatial genetic diversity of *Dysaphis plantaginea* in Europe, Morocco and North America (*N* = 582, clonal copies were excluded, 52 sites, 30 SSR markers). **a**. Map of allelic richness per site. **b**. Map of observed heterozygosity. Sites with a sample size below five individuals are not represented on this map. Each dot is a site (*i*.*e*., orchard). Red means high allelic richness, blue means low.

### Population structure and subdivision

The spatial genetic structures inferred for *D. plantaginea* with TESS, DAPC and STRUCTURE, and the respective *DIC, BIC* and *ΔK*, are shown in Figure 2 and supplementary material Figures S4 to S9. For each *K* value, CLUMPP analyses produced highly similar clustering patterns among the 10 repetitions (average *G* > 95%). We therefore only presented here the major modes.

**Figure 2.**
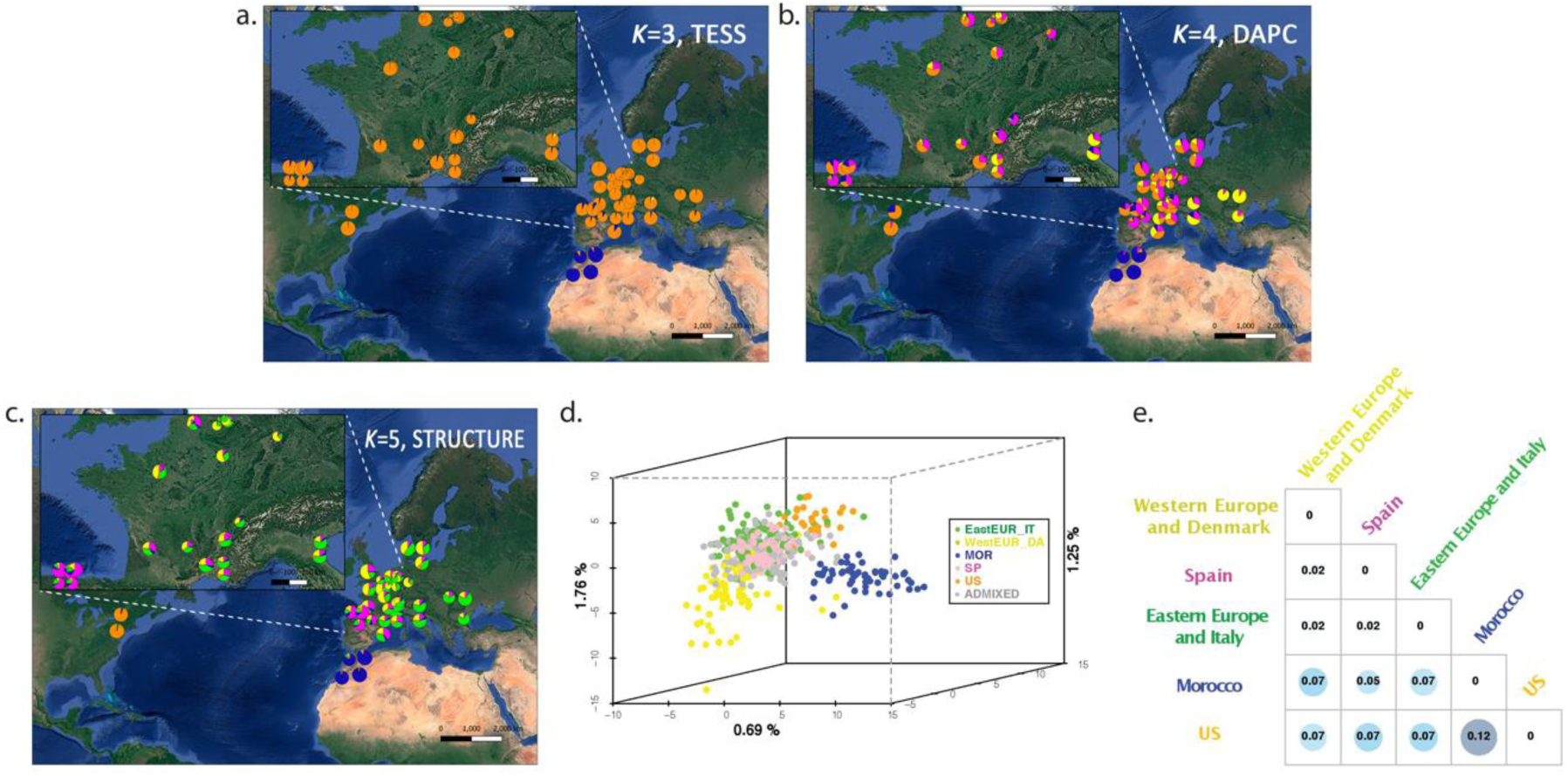
Spatial population structure and differentiation of the rosy apple aphid *Dysaphis plantaginea* in Europe, Morocco and North America (*N* = 582 individuals, 52 sites, 30 SSR markers). Population structure inferred with **a**. TESS (*K* = 3), **b**. Discriminant Analysis of Principal Components (*K* = 4), **c**. STRUCTURE (*K* = 5). Each figure includes a map with each pie chart representing the mean membership coefficients for each site. **d**. Principal component analysis including the 582 individuals used for running STRUCTURE analyses; each individual is colored according to its membership > 0.625 to one of the five *D. plantaginea* populations detected with STRUCTURE (individuals with a membership coefficient < 62.5% to a given cluster is referred as “admixed”). The size of each pie chart is equivalent to the number of individuals sampled at each site. **e**. Genetic differentiation (*F*_*ST*_) estimates among the five populations detected with STRUCTURE. The highest *F*_*ST*_ values are highlighted by circles (Morocco and the US has the highest value). US: United States of America; MOR: Morocco; SP: Spain; EasternEU_IT: Eastern Europe and Italian population (Bulgaria, Italy, Romania); WestEUR_DA: Western Europe and Danish population (France, Germany, Belgium, United Kingdom, and Denmark).

With TESS, increasing *K* above 3 did not reveal any additional cluster (Figure S4). For *K* = 3, TESS analyses revealed a clear partition between the Moroccan samples (blue), and other samples (European and North American, orange). An additional cluster was identified comprising only one Italian individual (yellow) (Figure 2a). With DAPC, increasing *K* above 4 did not reveal well-delimited new clusters, *i*.*e*., only individuals with multiple admixtures were assigned to the new clusters (Figure S5). For *K* = 4, DAPC identified three well-delimited clusters (Figure 2b): one in Eastern Europe (*i*.*e*., Bulgaria, Italy and Romania, in yellow), one in Morocco (blue), one in the USA (orange), plus another cluster that comprised the rest of the Western European (Spain, France, Belgium, the UK) and Danish samples (pink). With STRUCTURE, increasing *K* above 5 did not reveal additional clusters. For *K* = 5 (Figure 2c), samples were partitioned as follows: Morocco (blue), the USA (orange), Spain (pink), Eastern Europe and Italy (green), and Western Europe and Denmark (yellow).

We used the inferences from STRUCTURE in subsequent analyses because it revealed the finest population genetic structure. Genotypes were then assigned to a given population if their membership coefficient for that population exceeded 0.625 (Tables 1 and S1). We chose this threshold based on the bimodal distribution of cumulative coefficients inferred with STRUCTURE for *K* = 5 (Figure S10). A total of 175 admixed individuals (30% of the dataset) could not be assigned to any population and were not included in subsequent analyses. Most of the admixed individuals were located in Western and Northern Europe; the spatial distribution of the mean number of admixed individuals per site is represented in Figure S11.

**Table 1.**
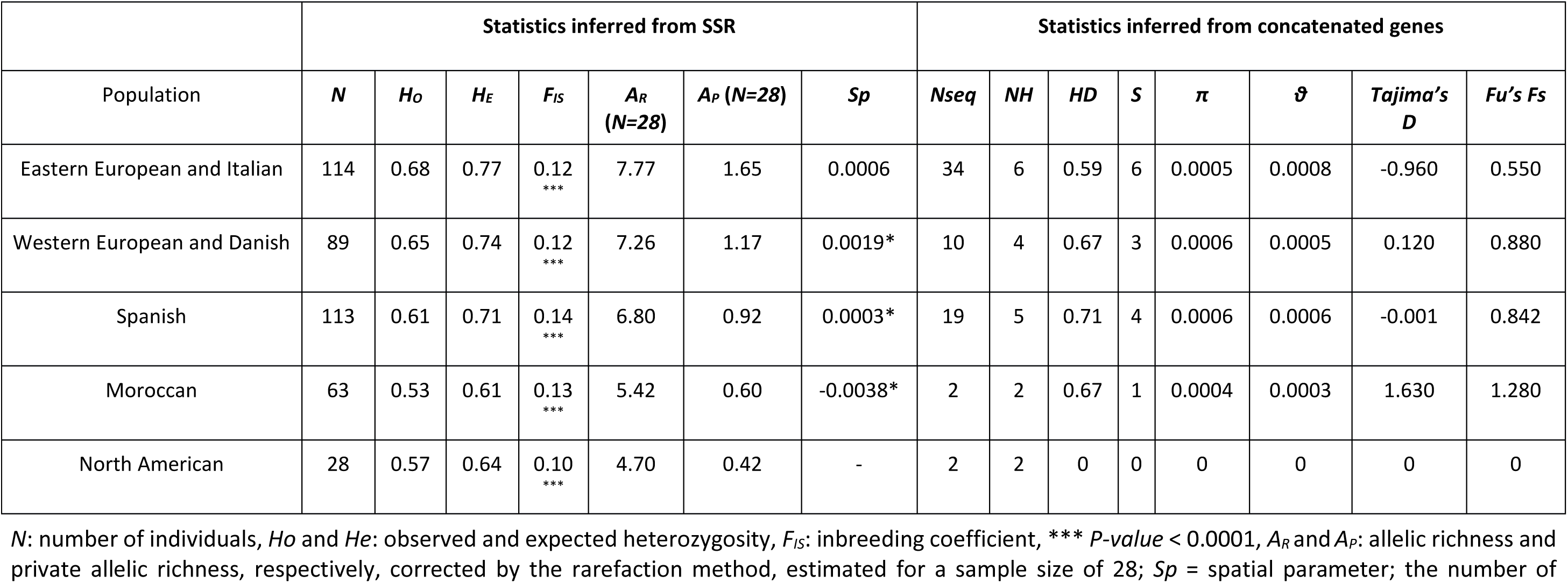
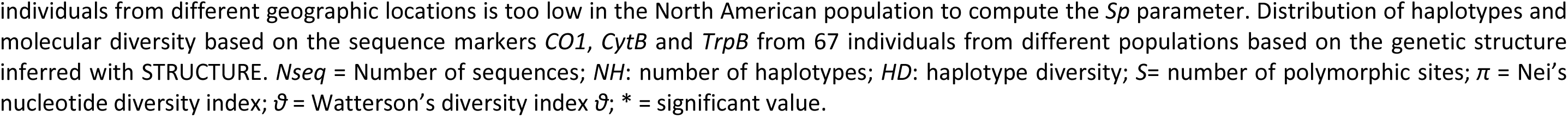
Summary statistics obtained for the five *Dysaphis plantaginea* populations (*i*.*e*., cluster including individuals with a membership coefficient > 62.5% to a given cluster) inferred with STRUCTURE (*K* = 5), based on 30 SSR markers and three concatenated genes (*CO1, Cytb* and *TrpB*, 67 *D. plantaginea* individuals), respectively (left and the right sections of the table).

| Population | Statistics inferred from SSR |  |  |  |  |  |  | Statistics inferred from concatenated genes |  |  |  |  |  |  |  |
| --- | --- | --- | --- | --- | --- | --- | --- | --- | --- | --- | --- | --- | --- | --- | --- |
| | $N$ | $H_o$ | $H_e$ | $F_{IS}$ | $A_R$<br>($N=28$) | $A_P$ ($N=28$) | $Sp$ | $N_{seq}$ | $NH$ | $HD$ | $S$ | $\pi$ | $\theta$ | Tajima's $D$ | Fu's $F_s$ |
| Eastern European and Italian | 114 | 0.68 | 0.77 | 0.12<br>*** | 7.77 | 1.65 | 0.0006 | 34 | 6 | 0.59 | 6 | 0.0005 | 0.0008 | -0.960 | 0.550 |
| Western European and Danish | 89 | 0.65 | 0.74 | 0.12<br>*** | 7.26 | 1.17 | 0.0019* | 10 | 4 | 0.67 | 3 | 0.0006 | 0.0005 | 0.120 | 0.880 |
| Spanish | 113 | 0.61 | 0.71 | 0.14<br>*** | 6.80 | 0.92 | 0.0003* | 19 | 5 | 0.71 | 4 | 0.0006 | 0.0006 | -0.001 | 0.842 |
| Moroccan | 63 | 0.53 | 0.61 | 0.13<br>*** | 5.42 | 0.60 | -0.0038* | 2 | 2 | 0.67 | 1 | 0.0004 | 0.0003 | 1.630 | 1.280 |
| North American | 28 | 0.57 | 0.64 | 0.10<br>*** | 4.70 | 0.42 | - | 2 | 2 | 0 | 0 | 0 | 0 | 0 | 0 |

$N$ : number of individuals, $H_o$ and $H_e$ : observed and expected heterozygosity, $F_{IS}$ : inbreeding coefficient, \*\*\* $P$ -value < 0.0001, $A_R$ and $A_P$ : allelic richness and private allelic richness, respectively, corrected by the rarefaction method, estimated for a sample size of 28; $Sp$ = spatial parameter; the number of
individuals from different geographic locations is too low in the North American population to compute the $S_p$ parameter. Distribution of haplotypes and molecular diversity based on the sequence markers *CO1*, *CytB* and *TrpB* from 67 individuals from different populations based on the genetic structure inferred with STRUCTURE. $N_{seq}$ = Number of sequences; $NH$ : number of haplotypes; $HD$ : haplotype diversity; $S$ = number of polymorphic sites; $\pi$ = Nei's nucleotide diversity index; $\vartheta$ = Watterson's diversity index $\vartheta$ ; \* = significant value.

We further visualized and quantified the genetic differentiation among populations with a principal component analysis (3D-PCA Figures 2d and S12) and *F*_*ST*_ estimates (Figure 2e), respectively. Among all populations, *F*_*ST*_ values were low but all significant (Figure 2e). Pairwise genetic differentiation was the lowest between the European populations (*F*_*ST*_ = 0.02, *P-value* < 0.0001, Figure 2e); the Moroccan and the USA populations were the most differentiated (Figure 2e). The PCA showed similar genetic relationships among the five populations (Figure 2d).

### Genetic diversity, bottleneck and range expansion

The mean expected heterozygosity was relatively high (average = 0.74 with values ranging from 0.55 to 0.75, Table S8). The mean *F*_*ST*_ across all loci was low (mean *F*_*ST*_ = 0.08, range: -0.006 - 0.5) but significant for all pairs of sites (*P-value* < 0.001). Allelic richness and private allelic richness were significantly different among the five populations, except between the North American and Moroccan populations for allelic richness, and among the Spanish, Moroccan and Eastern European populations for private allelic richness (Table S9). The North American and Moroccan populations showed significantly lower levels of allelic richness and private allelic diversity than the European populations (*i*.*e*., Spanish, Eastern and Western European populations; Tables 1 and S9). In Europe, the Eastern European/Italian population had the highest level of allelic richness and private allelic diversity, followed by the Western European/Danish population, and lastly the Spanish population.

We also tested whether a strong and recent bottleneck occurred for each population. BOTTLENECK analyses showed no significant deviation from the mutation-drift equilibrium in any of the populations (Table S10). Tajima’s *D* and Fu’s *F*_*S*_ statistics did not reveal any signature of demographic range expansion either (Table 1).

### Isolation-by-Distance (IBD)

A significant but weak IBD pattern was observed for the rosy apple aphid, using the 52 sampling sites (*r*^*2*^=0.057, *P-value* ≤ 0.05), or, excluding the USA sampling sites (*i*.*e*., 50 sites, *r*^*2*^=0.041, *P-value* ≤0.05), or, using only the European populations (*i*.*e*., 45 sites, *r*^*2*^=0.047, *P-value* ≤ 0.05 (Figure S13). The *Sp* statistic can be used to quantify spatial structure and is useful for comparing populations and/or species. Low *Sp* values are associated with greater dispersal capacities and/or effective population sizes. Here, *Sp* values were extremely low (close to 0) and were only significant for the Moroccan and the Western European/Danish populations (Table 1). These results suggest that *D. plantaginea* has high dispersal capacities and/or large effective population sizes.

### Inference of the divergence and demographic history of the rosy apple aphid

First, we reconstructed the divergence and demographic history of the rosy apple aphid in Europe (*i*.*e*., including only the Spanish, Eastern European/Italian and Western European/Danish populations). We defined 12 scenarios assuming different divergence histories of the Eastern European/Italian, Western European/Danish and Spanish populations (Figure S2). The 12 scenarios were tested with and without gene flow among populations. We therefore ended up comparing 24 scenarios. Classification votes from the first round were the highest ten times out of ten for the group of scenarios that assumed gene flow among the three populations (295 votes out of the 500 RF trees, posterior probability *P =* 0.61, prior error rate = 3.04%, Table S12, Figure S14). Projection of the reference table datasets and the observed dataset on a single axis showed that the observed data fell within the distribution of the simulated summary statistics of the group of scenarios that assumed gene flow among the three populations, suggesting this analysis had the power to discriminate between the two groups of scenarios and to identify the most likely scenario (Figure S14). The second round of ABC inferences testing the sequence of colonization of the European populations requires caution in interpretation as prior error rates were high and posterior probabilities low (Figure S14 and Table S15).

We then investigated the colonization history of the rosy apple aphid outside of Europe, *i*.*e*., of the Moroccan and North American populations. Given the lack of power to discriminate between different scenarios of colonization sequence of the rosy apple aphid in Europe, and the weak genetic differentiation among the three European populations (mean *F*_*ST*_ = 0.02, Figure 2e), we merged the three European populations into a single European population (*N* = 316) for this analysis. We then defined six scenarios of sequence of colonization starting either from Europe or Morocco (Figure S2). We excluded the hypothesis that the rosy apple aphid originated in North America. Indeed, historical records show that the introduction of the rosy apple aphid was very recent in America (*ca*. 1890s) (Foottit et al., 2006). Furthermore, the North American population had the lowest levels of private allelic diversity and allelic richness, and the North American samples clustered with the European samples in the DAPC and TESS analyses. For each of the six scenarios, five scenarios of gene flow among populations were tested: no gene flow, gene flow among all populations and gene flow between each population pair (*i*.*e*., Europe/Morocco, Europe/the USA, and Morocco/the USA). We simulated these specific models of gene flow among specific pairs of populations for the ABC-RF as we observed variable admixture levels among populations (Figure 2c). In total, 30 scenarios were compared (six colonization sequences x five gene flow modes, Figure S2). ABC-RF analyses showed relatively high support for scenarios assuming gene flow (ABC-RF round 1, 10 out of the 10 replicates for the groups of scenarios assuming gene flow, 337 votes out of the 500 RF trees, posterior probability *P =* 0.65, prior error rate = 6.55%, Table S14, Figure S15). Projection of the reference table datasets and the observed dataset on a single axis showed that the observed data fell within the distribution of the simulated summary statistics of the group assuming gene flow, suggesting the occurrence of gene flow (Figure S15). However, although the observed dataset fell within the distribution of simulated summary statistics (Figure S15), we lacked the power to infer the sequence of colonization of the rosy apple aphid outside of Europe (*i*.*e*., posterior probability *P =* 0.65, and high prior error rate = 73.8%, Table S15).

Altogether, ABC-RF inferences supported the occurrence of gene flow during the colonization history of *D. plantaginea*. However, ABC-RF did not allow to determine the sequence of colonization of Europe, North America and Morroco by the rosy apple aphid.

### 16S rRNA amplicon sequencing

After sequence filtering using the FROGS pipeline, high-throughput sequencing of 16S rRNA bacterial genes from 186 aphids resulted in 5.7 M sequencing reads with an average of 30,800 reads per aphid sample. We found an extremely low bacterial diversity in *D. plantaginea*. The 5.7 M sequencing reads were clustered into 18 OTUs (Figure S16, Table S1). Seven OTUs were assigned to *B. aphidicola* and made up 97.8% of the sequencing reads, 92 % of the reads were assigned to a single *B. aphidicola* OTU, which was found associated with all *D. plantaginea* individuals, the remaining *B. aphidicola* OTUs were found associated with “outgroups”. The remaining reads were mainly assigned to two known aphid endosymbionts, *Serratia symbiotica* and *Regiella insecticola*, which were found in eight and three aphid specimens, respectively. The three aphids hosting *Regiella* belonged to a population of *M. pyraria* collected in Switzerland, while aphids hosting *Serratia* were found in distantly related populations including *D. plantaginea* collected on *M. domestica* from various apple orchards in France, and *P. communis* in Iran (Table S1). These results highlight the extremely limited diversity of symbionts in the rosy apple aphid across a large geographical scale.

## Discussion

We investigated the demographic history of a major fruit tree pest, the rosy apple aphid, in the regions where it impacts the most cultivated apple orchards (*i*.*e*., Europe, North Africa and North America). Using multiple approaches, we showed that the colonization of Europe by the rosy apple aphid is likely recent, was not accompanied by strong bottlenecks and involved gene flow between and within populations. The high level of gene flow among populations and within populations was supported by the weak spatial genetic structure observed across Europe, and coalescent-based simulations combined with ABC-RF. We also found that *D. plantaginea* rarely hosts endosymbiotic bacteria other than their primary symbiont, *B. aphidicola* in North America and Europe. Our results provide further understanding of the evolutionary processes at play during pest range expansion.

### Colonization of apple trees by the rosy apple aphid is recent, and did not involve drastic bottlenecks

After removing clonal copies from our analyses, we detected with STRUCTURE five main panmictic populations of the rosy apple aphid: three in Europe, one in Morocco and one in the USA. The genetic diversity for each population was within the same range as other aphid species such as *B. helichrysi* (Popkin et al., 2017), *Eriosoma lanigerum* (Zhou et al., 2015) and *M. persicae nicotianae* (Zepeda-Paulo et al., 2010). However, genetic diversity (private alleles and allelic richness) in *D. plantaginea* was higher in Europe and lower in the USA and Morocco. More generally, patterns of genetic structure and diversity suggest different demographic histories for the European, Moroccan and American populations of *D. plantaginea*.

Individuals from the European populations had partial memberships to multiple clusters, with similar membership coefficients for most individuals. This pattern of high admixture along a spatial transect might reflect a continuous gradation in allele frequencies (*i*.*e*., a cline) across regions that cannot be detected by the methods used in this study. Indeed, a major limitation of all clustering approaches is the risk of inferring artefactual discrete groups in populations where genetic diversity is distributed continuously. DAPC and STRUCTURE are not immune to this bias and may erroneously identify clusters within a cline (Jombart et al., 2010). TESS is more sensitive to allelic gradient; this program includes a decay of the correlation between membership coefficients and distance within clusters (Chen et al., 2007). In the presence of clines and with evenly distributed sampling, TESS may detect fewer clusters than STRUCTURE (Durand, Chen, & François, 2009). The larger admixed clusters found with DAPC and STRUCTURE may therefore reflect a cline of allele frequency across Europe for the rosy apple aphid. Allele frequency clines can result from admixture between genetically distinct populations (Currat & Excoffier, 2005; Menozzi, Piazza, & Cavalli-Sforza, 1978) and/or from subsequent founder events during range expansion (Barbujani, Sokal, and Oden 1995; Fix 1997; Currat and Excoffier 2005). Genetic diversity is also expected to decrease along the expanding range (François, Blum, Jakobsson, & Rosenberg, 2008; Prugnolle, Manica, & Balloux, 2005). Founder events associated with a recent range expansion may have resulted in the decreasing east-west gradient of genetic diversity and the large number of admixed individuals observed in Europe. It is therefore possible that the European populations of the rosy apple aphid underwent a recent expansion. Note that we did not detect any signature of range expansion with the three-markers dataset (*CO1, cytB* and *trpB*). This lack of signature of range expansion may be due to limited number of samples used in our test (at least for the Moroccan and North American populations), but also that the range expansion is so recent that we cannot catch its footprint with our sequence markers. Several preliminary ABC-RF tests of range expansion of the rosy apple aphid in Europe that failed (*i*.*e*., very low posterior probabilities and prior error rates, data not shown) suggest that the second hypothesis is possible.

The Moroccan and North American populations displayed a different pattern of genetic differentiation and diversity compared to the European populations, suggesting an even more recent colonization history. The two populations were well circumscribed and had the highest level of genetic differentiation from the European populations, the lowest genetic diversity and the lowest number of private alleles. In the TESS analysis, the North American samples did not cluster separately from the European samples, and in the DAPC analysis, samples from North America and Western Europe clustered together, suggesting that the North American population originated recently from Europe. This agrees with the earliest record of *D. plantaginea* in the Eastern US dating back to 1890 (Foottit et al., 2006). Since then, there have probably been multiple introductions into the USA that prevented genetic differentiation from Europe. Similar scenarios have been described for the tobacco aphid, which was introduced into America from different European gene pools (Zepeda-Paulo et al., 2010), and the leaf-curl plum aphid *B. helichrysi* (Piffaretti et al., 2013), for which population genetic tools showed very little differentiation between European and North American populations. As for the Moroccan population, we lacked the power to infer its origin with the ABC-RF method. However, the significantly lower genetic diversity and number of private alleles in this population compared with that of Europe, the high level of admixture and the close genetic relationship with the Spanish population, suggest that the Moroccan population resulted from a recent colonization event from Southern Europe. Nevertheless, the history of the rosy apple aphid in North Africa deserves further investigation requiring additional sampling in this region. Altogether, the significantly lower genetic diversity observed in the Moroccan and North American populations suggest that these originated recently through founder events. Indeed, we did not find any evidence that these two populations underwent a recent strong bottleneck despite having significantly lower diversity in both SSR and sequence markers. Thus, the founder effect underlying the colonization of North America and Morocco may involve genetic drift in small populations rather than severe bottlenecks at the introduction event. Genetic diversity was also found to be higher in native populations within their native range in the tobacco aphid (Zepeda-Paulo et al., 2010) and the Russian wheat aphid (Zhou et al., 2015).

Altogether, a recent range expansion of the rosy apple aphid on its cultivated apple host is a plausible explanation of the observed spatial genetic structure and diversity. Rapid range expansion has also been described in the Russian wheat aphid (Zhang et al., 2014). Unfortunately, the ABC-RF method was not powerful enough to disentangle the different scenarios of colonization of the rosy apple aphid. However, the ABC-RF method was powerful enough to identify the occurrence of gene flow within and outside Europe. Another recent study on the main invasion routes of *D. suzukii* also reported a moderate level of confidence in model choice (*i*.*e*., low posterior probabilities between 0.50 to 0.63) and high prior error rates (ranging from 0.30 to 0.40; using ABC-RF (Fraimout et al., 2017)). The absence of samples from key places during the species colonization and individuals of the species ancestral group have been discussed to impact in the resolution of the colonization history analysis using ABC approaches (Lombaert et al., 2014). This might account for the lack of support for some of our colonization scenarios using ABC-RF. Additional information on the mutation rates of the newly developed SSR in the present study may also improve support to some of our colonization scenarios. The mutation rate of transcribed and untranscribed SSR was indeed successfully used to reconstruct the main migration routes of *S. gregaria* to Africa using ABC-RF (Chapuis et al., 2020).

### Colonization with gene flow, likely driven by humans

We found that the expansion of the rosy apple aphid involved several populations with high genetic diversity each, and a high extent of historical gene flow for *D. plantaginea* across Europe, North America and Morocco. Population structure analyses indicated that 30 % of the individuals was a product of recent admixture. Once recently admixed individuals removed, ABC-RF inferences strongly support scenarios with bidirectional gene flow among populations. Both population structure and coalescent-based method inferences therefore indicate the occurrence of recent and ancient gene flow among *D. plantaginea* populations. In addition, the *Sp* parameter estimates revealed large extent of historical gene flow within population. We used the *Sp* parameter estimates to compare the dispersal capacities of *D. plantaginea* with existing estimates in plants (Vekemans & Hardy, 2004). The rosy apple aphid showed dispersal capacities equivalent to or even higher than that of wind-dispersed trees. The rosy apple aphid can therefore spread over long distance, as suggested previously (Guillemaud et al., 2011), and as frequently found in aphids (Loxdale et al., 1993). However, despite its high dispersal capacities and the large amount of gene flow among and within populations, we can still observe a significant (but weak) spatial genetic structure across its distribution. The observed subtle spatial genetic structure may be associated with agricultural practices. The interchange of apple materials (*i*.*e*., cultivars in the form of scions and/or trees) is nowadays, and probably has been historically, frequent and a potential way of moving the rosy apple aphid in the form of overwintering eggs among regions where apple is cultivated. This hypothesis agrees with the differentiated population of *D. plantaginea* observed in Asturias (Northwestern Spain). This region is known for the main utilization of the apple for cider production and is based on local cider cultivars (Tardío, Arnal, & Lázaro, 2020). Most of our samples from Spain are from Asturias, except the Catalan samples that clustered with the European ones. The exchange of apple material within Asturias, and within other regions in Europe, may have occurred, implying higher risk of aphid movement within regions than between regions. Of course, this is not the only explanation of the spatial genetic structure observed, physical barriers (*i*.*e*., Atlantic Ocean) could also explain the higher genetic differentiation of the Moroccan and North American populations.

### Putative center(s) of origin of the rosy apple aphid

The geographical origin of *D. plantaginea* remains unresolved. Our phylogenetic analyses confirmed previous relationships between the rosy apple aphid and other aphids (Bašilova & Rakauskas, 2012; Rebijith et al., 2017). However, estimates of the divergence time of *D. plantaginea* from closely related species is now required, with denser sampling of the *Dysaphis* genus across several regions in Eurasia and molecular dating analyses.

Genetic diversity estimates from SSR markers suggest that the source population came from Eastern Europe, but it may also have originated even further east in Central Asia where its fruit tree host was originally domesticated (Cornille et al., 2014; Harris et al., 2002). Our failure to collect *D. plantaginea* samples in this area, despite our attempts in China and Kazakhstan, prevents us from testing this scenario. However, note that, while *D. plantaginea* has been recorded in Central Asia according to several faunistic surveys (Kadyrbekov, 2002), it is hard to find in these regions. Furthermore, a lack of records of this species in the Global Biodiversity Information Facility (GBIF, 2020) suggests *D. plantaginea* is uncommon in this area.

Alternatively, *D. plantaginea* may have originated in the Caucasus or Asia Minor, maybe through a host jump from Pyrus to Malus. *Dysaphis plantaginea, D. radicola, D. devecta, D. brancoi, D. anthrisci, D. chaerophylli* are *Dysaphis* species reported to feed on the cultivated apple, *M. domestica*, as their primary host (Blommers et al., 2004; Stekolshchikov, 2006), but many aphid species also feed on pears, including, *D. reaumuri* Mordvilko and *D. pyri* Boyer de Fonscolombe (Barbagallo, Cocuzza, Cravedi, & Komazaki, 2007). The *Pyrus* genus is known to have diverged a long time ago from the genus *Malus* probably in the Caucasus (Celton et al., 2009; Xiang et al., 2017). Therefore, the ancestral group from which *D. plantaginea* diverged might have had a *Pyrus* species as host. Analysis of samples from these regions is required to test this hypothesis.

### Low endosymbiont bacterial diversity associated with *D. plantaginea*

Our results showed that bacterial diversity was strikingly low in *D. plantaginea* across Europe and North America. As expected, *B. aphidicola* was the predominant bacterial species (97% of our reads). We also distinguished different variants of *B. aphidicola* in the Iranian *D. plantaginea* samples which fits with the genetic differentiation observed in the corresponding aphid hosts. At least nine secondary endosymbionts have been reported among aphid species (reviewed in Zytynska & Weisser, 2016), including *Arsenophonus* Gherma, *Hamiltonella defensa* Moran, *Regiella insecticola, Rickettsia* Da Rocha-Lima, *Rickettsiela* Drobne, *Serratia symbiotica, Spiroplasma* Saglio and *Wolbachia* Hertig & Wolbach, and *Fukatsuia symbiotica*. These bacteria have been reported to be aphid facultative symbiont, potential pathogens, or plant-associated microbiota (Gauthier, Outreman, Mieuzet, & Simon, 2015). Here, we detected the presence of secondary symbionts only in a few samples. The *Serratia* bacteria was the only secondary symbiont identified and it was found in only eight samples of *D. plantaginea*, collected on *Pyrus communis* in Iran, France and Spain on *M. domestica. Regiella* symbiont was found in three samples from another aphid species, *M. pyraria* collected on *M. domestica*. At least seven endosymbionts (*H. defensa, R. insecticola, Rickettsia* sp., *Rickettsiella* sp., *S. symbiotica, Spiroplasma* sp. and *Wolbachia* sp.) have been reported across a narrower spatial distribution in the model species *Acyrthosiphon pisum* and several other well-studied aphid species (Zytynska & Weisser, 2016). By contrast, another study (Henry, Maiden, Ferrari, & Godfray, 2015) found that neither *D. plantaginea* nor other *Dysaphis* species hosted any secondary symbionts, although it relied on a small number of individuals. This low endosymbiont bacterial diversity in *D. plantaginea* shows that its likely fast expansion is not the result of an association with different mutualistic endosymbionts.

### Concluding remarks

This study has demonstrated that the colonization of a major fruit tree aphid pest occurred without a strong bottleneck, maintaining high genetic diversity, and generated differentiated populations exchanging gene flow, with an isolation-by-distance pattern. The lack of demographic changes in the populations of *D. plantaginea* in Europe, except for the Belgian populations, indicates that seasonal selective pressures, such as insecticide application, have little impact on the genetic diversity of the species. These results may have implications in control and management of *D. plantaginea*, but further studies are needed to fully understand how selective pressures have impacted *D. plantaginea* adaptation. In addition, the use of other genetic markers, such as SNPs, promises to make great strides to elucidate the demographic history of the rosy apple aphid. Finally, the origin of the rosy apple aphid is still unknown. Our results suggest it may have originated in Eastern Europe, the Caucasus or Asia Minor. However, the domestication of the rosy apple aphid primary host (the cultivated apple) in this region remains unknown (Cornille et al., 2019, 2014; Spengler, 2019). Further investigations on the history of apple domestication, additional sampling of *D. plantaginea* in the Caucasus or Asia Minor, and sampling related aphid species are required to better understand the origin of this major fruit tree pest.

## Data accessibility

Data are available online: https://zenodo.org/record/4537710#.YCaeomMo95c

## Supplementary material

https://zenodo.org/record/4537710#.YCaeomMo95c

## Acknowledgements

This research was funded by the laboratoire d’Excellence Biodiversity Agrosystem Society and Climate BASC (grant « Emergence POMPUCEDOM ») and Systematic Research funding, ATIP-Avenir program and the *Institut Diversité Ecologie et Evolution du Vivant* (IDEEV). AR and TU were supported by a grant of the Romanian Ministry of Education and Research, CCCDI - UEFISCDI, project number 384 PED-PN-III-P2-2.1-PED-2019-4924. We thank the Plateforme de Génotypage GENTYANE INRAE UMR 1095 for assistance in genotyping, and especially the platform leader Charles Poncet. We thank the informatics team at the GQE-Le Moulon, in particular Adrien Falce, Benoit Johannet and Olivier Langella. We are grateful to the Migale bioinformatics Facility (MIGALE, INRAE, 2020, doi: 10.15454/1.5572390655343293E12) for providing help and/or computing and/or storage resources. Version 3 of this preprint has been peer-reviewed and recommended by Peer Community In Evolutionary Biology (https://doi.org/10.24072/pci.evolbiol.100134).

## Authors ‘contributions

AC, EJ, PD, LJ conceived and designed the experiments; AC, EJ, PD, LJ, TG, MH obtained funding; AC, AR, AA, TB, GA, FP, AH, SKJ, RA, SS, LS, EG, LT, FG, KM, YK, ARom, TD, IZ, OS, RJM, AA, BEH, EJ, HZ, IB, MR, HT, CM, BB, AD sampled the material; AR, CR, AV, OG, AC, MA, LM, ML, LB, SGOV performed the molecular work; SGOV, AC, OG, MG, LB, EJ, GD analyzed the data. All co-authors discussed the results. The manuscript was written by SGOV, AC, EJ, OG with critical inputs from other co-authors.

## Conflict of interest disclosure

The authors of this preprint declare that they have no financial conflict of interest with the content of this article. Tatiana Giraud is a member of the managing board of PCI Evolutionary Biology. Tatiana Giraud, Amandine Cornille, Emmanuelle Jousselin, and Jean-Christophe Simon are PCI Evolutionary Ecology Biology recommenders.

## Notes

https://zenodo.org/record/4537710#.YCaeomMo95c

### Competing Interest Statement

The authors of this preprint declare that they have no financial conflict of interest with the content of this article. Amandine Cornille, Tatiana Giraud, Emmanuelle Jousselin, and Jean-Christophe Simon are PCI Evolutionary Ecology Biology recommenders. Tatiana Giraud is a member of the managing board of PCI Evolutionary Biology.

### Summary of Updates

The manuscript has been revised according to PCI Evol Biol recommenders. The authors thank the reviewers for their very constructive suggestions.

