## Supporting information for "Large-scale geography survey provides insights into the colonization history of a major aphid pest on its cultivated apple host in Europe, North America and North Africa"

**Table S1.** **Information for each sample of *Dysaphis plantaginea* used in this study (*N* = 715).** For each sample, identification number, geographic origin, plant host, sampler identification, and its respective use for each dataset (SSR, 16S rRNA metabarcoding and/or three sequence markers, *i.e.*, CO1, CytB, TrpB) are provided. Membership coefficients for *K*=5 inferred with STRUCTURE using 30 microsatellite markers, and OTU assignations obtained with FROGS using the 16S rRNA marker are also provided.

See table_S1.xlsx

**Figure S1.** Map of the 52 sampling sites of *Dysaphis plantaginea* used in this study (*N* = 667 individuals). Each dot represented one sampling site (*i.e.*, orchard).

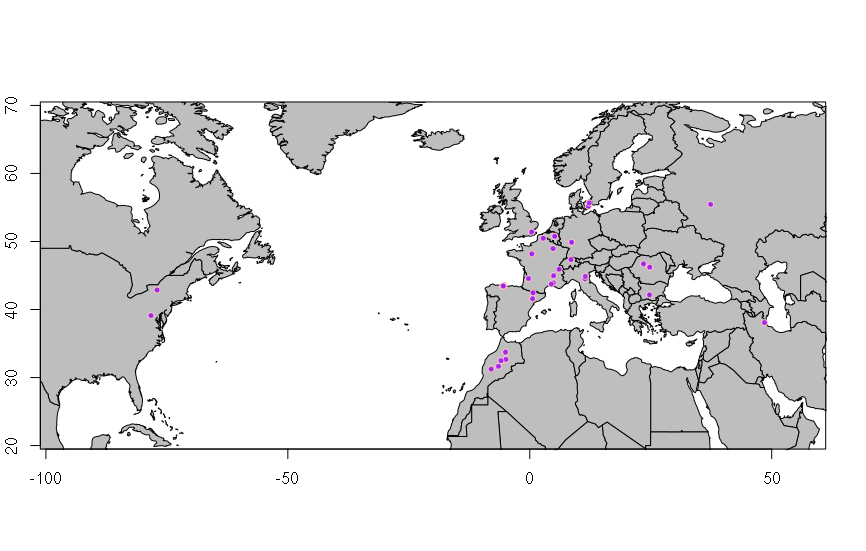

**Table S2.** Information on the sequence markers (*CO1, CytB, and TrpB*) used in this study on *Dysaphis plantaginea* samples, including sequences of the primers (Forward and Reverse), the expected size of the fragments (in base pairs), the temperatures of hybridization to achieve PCRs, and the source of the development of the markers.

| **Name** | **Gen** | **Sequence** | **Size**  **(bp)** | **PCR** | **Reference** |
| --- | --- | --- | --- | --- | --- |
| Cytochrome c oxidase I | CO1 | **F:** ATTCAACCAATCATAAAGATATTGG  **R:** TAAACTTCTGGATGTCCAAAAAATCA | 658 | 48°C, 1 min | (Sheffield, Hebert, Kevan, & Packer, 2009) |
| Cytochrome B | CytB | **F:** GATGATGAAATTTTGGATC  **R:** ATTACACCTCCTAATTTATTAGGAAT | 750 | 48°C, 1 min | (Jermiin, Olsen, Mengerson, & Easteal, 1997) |
| Tryptophan synthase beta | trpb | **F:** GCATGAAATTAATGGGTGCAG  **R:** ATCCTGCTGAAATKGACCAA | 450 | 55°C, 1 min | (Jousselin, Desdevises, & Coeur d’acier, 2009) |

**Table S3.** McDonald and Kreitman’s test computed with the Standard and Generalized McDonald-Kreitman test website (Egea, Casillas, & Barbadilla, 2008) for the sequence markers COI, CytB, and TrpB, and using *Aphis pomi* species as an external group.

|  |  |  |  |  |  |  |  | **J&C** |
| --- | --- | --- | --- | --- | --- | --- | --- | --- |
|  |  | **PM** | **DIV** | **Total** | **NI** | ***α*** | ***χ2*** | ***P-value*** |
|  | **Neutral** | 7 | 0.00 | 7 | null | null | null | null |
| **CO1*** | **Non-neutral** | 176 | 0.00 | 176 |  |  |  |  |
|  | **Total** | 183 | 0.00 | 183 |  |  |  |  |
|  | **Neutral** | 90 | 3 | 93 | null | null | 2.175 | 0.140 |
| **TrpB** | **Non-neutral** | 64 | 0 | 64 |  |  |  |  |
|  | **Total** | 154 | 3 | 157 |  |  |  |  |
|  | **Neutral** | 65 | 0 | 65 | 0.000 | 1.000 | 1.042 | 0.307 |
| **CytB** | **Non-neutral** | 62 | 1 | 63 |  |  |  |  |
|  | **Total** | 127 | 1 | 128 |  |  |  |  |

PM: polymorphism; DIV = divergence; Neutrality Index Jukes & Cantor = NI; Proportion of adaptive substitutions (α) Jukes & Cantor; χ2 value Jukes & Cantor = χ2 value; *P-value* < 0.05.

**Table S4.** Results of the Ewens-Watterson test for each of the 30 SSR markers (sorted by panels, see Text S3) for *Dysaphis plantaginea* computed with the POPGENE software (Yeh et al., 1997). These statistics were calculated using 100 simulated samples.

| **Panel** | **Locus** | ***N*** | ***k*** | **Obs.F** | **Min.F** | **Max.F** | **Mean^*^** | **SE^*^** | **L95^*^** | **U95^*^** |
| --- | --- | --- | --- | --- | --- | --- | --- | --- | --- | --- |
| **Alexa fluor** | **SSR16** | 1322 | 35 | 0.255 | 0.0286 | 0.9499 | 0.1315 | 0.0021 | 0.0749 | 0.2454 |
|  | **SSR19** | 1294 | 18 | 0.2061 | 0.0556 | 0.9741 | 0.2507 | 0.0095 | 0.1315 | 0.521 |
|  | **SSR2** | 1292 | 37 | 0.0837 | 0.027 | 0.9458 | 0.1258 | 0.0018 | 0.0724 | 0.2322 |
|  | **SSR3** | 1310 | 28 | 0.0834 | 0.0357 | 0.9596 | 0.1616 | 0.0036 | 0.089 | 0.3242 |
|  | **SSR49** | 998 | 12 | 0.2516 | 0.0833 | 0.9782 | 0.3505 | 0.0194 | 0.1714 | 0.7087 |
|  | **SSR5** | 1328 | 10 | 0.542 | 0.1 | 0.9865 | 0.4076 | 0.024 | 0.1888 | 0.7906 |
|  | **SSR6** | 1300 | 18 | 0.246 | 0.0556 | 0.9742 | 0.2517 | 0.0101 | 0.1314 | 0.5226 |
|  | **SSR8** | 1328 | 12 | 0.4636 | 0.0833 | 0.9836 | 0.357 | 0.0195 | 0.1784 | 0.7161 |
| **ATTO 550** | **SSR18** | 1300 | 9 | 0.3519 | 0.1111 | 0.9878 | 0.4351 | 0.027 | 0.2156 | 0.8367 |
|  | **SSR23** | 1146 | 8 | 0.3503 | 0.125 | 0.9879 | 0.4603 | 0.0292 | 0.2281 | 0.8679 |
|  | **SSR26** | 1300 | 19 | 0.2817 | 0.0526 | 0.9727 | 0.2373 | 0.0081 | 0.1227 | 0.4743 |
|  | **SSR27** | 1310 | 20 | 0.3057 | 0.05 | 0.9714 | 0.2342 | 0.0087 | 0.1225 | 0.4689 |
|  | **SSR28** | 1318 | 10 | 0.3469 | 0.1 | 0.9864 | 0.4033 | 0.0244 | 0.1961 | 0.7893 |
|  | **SSR32** | 1268 | 45 | 0.2075 | 0.0222 | 0.933 | 0.0998 | 0.001 | 0.0585 | 0.1709 |
|  | **SSR51** | 1160 | 31 | 0.1062 | 0.0323 | 0.9496 | 0.1449 | 0.0029 | 0.0819 | 0.2855 |
| **ATTO 565** | **L4** | 1232 | 20 | 0.3849 | 0.05 | 0.9696 | 0.2258 | 0.0082 | 0.1168 | 0.4594 |
|  | **SSR17** | 1296 | 25 | 0.1323 | 0.04 | 0.9636 | 0.1834 | 0.0052 | 0.1022 | 0.3832 |
|  | **SSR21** | 1168 | 11 | 0.3291 | 0.0909 | 0.983 | 0.3809 | 0.0225 | 0.1782 | 0.7519 |
|  | **SSR38** | 1286 | 25 | 0.1662 | 0.04 | 0.9634 | 0.1842 | 0.0045 | 0.0999 | 0.3613 |
|  | **SSR4** | 1318 | 39 | 0.146 | 0.0256 | 0.944 | 0.1166 | 0.0014 | 0.07 | 0.2149 |
|  | **SSR40** | 1302 | 24 | 0.1874 | 0.0417 | 0.9653 | 0.1935 | 0.0057 | 0.1022 | 0.3972 |
|  | **SSR7** | 1270 | 11 | 0.4099 | 0.0909 | 0.9844 | 0.3817 | 0.0226 | 0.185 | 0.7414 |
| **HEX** | **SSR1** | 1290 | 41 | 0.0874 | 0.0244 | 0.9399 | 0.1092 | 0.0014 | 0.0637 | 0.2088 |
|  | **SSR11** | 1326 | 14 | 0.4268 | 0.0714 | 0.9806 | 0.3103 | 0.0143 | 0.1521 | 0.6184 |
|  | **SSR13** | 1322 | 8 | 0.3454 | 0.125 | 0.9895 | 0.4676 | 0.0297 | 0.2303 | 0.8575 |
|  | **SSR15** | 1304 | 50 | 0.2589 | 0.02 | 0.9277 | 0.0879 | 0.0007 | 0.0547 | 0.1558 |
|  | **SSR22** | 1326 | 16 | 0.2792 | 0.0625 | 0.9776 | 0.2832 | 0.0124 | 0.1468 | 0.5763 |
|  | **SSR31** | 1330 | 13 | 0.1452 | 0.0769 | 0.9821 | 0.3342 | 0.0192 | 0.1657 | 0.6945 |
|  | **SSR46** | 1256 | 16 | 0.1693 | 0.0625 | 0.9764 | 0.2815 | 0.0121 | 0.1406 | 0.579 |
|  | **SSR9** | 1310 | 15 | 0.2737 | 0.0667 | 0.9789 | 0.2934 | 0.0134 | 0.1492 | 0.598 |

*N*: sample size; *k*: number of alleles; Obs.F: Observed allelic frequency; Min.F: Minimum allelic frequency; Max.F: Maximum allelic frequency; Mean: mean allelic frequency; SE: standard error of the mean; L95 and U95: Lower and Upper 95% confidence interval, respectively.

**Table S5.** Details of primer sequence (Forward and Reverse), repeat motif, allelic range size, annealing temperature, and population genetic summary statistics for each of the 30 microsatellite markers used for reconstructing population structure, genetic diversity and colonization history of *Dysaphis plantaginea* (statistics were calculated before removal of the clones, *i.e.*, including 667 individuals).

| **Dye** | **Locus** | **Primer sequences (5’-3’)** | **Repeat** | **Range** | **T°** | **Panel** | **Alleles** | ***H_O_*** | ***H_E_*** | ***F_IS_*** | ***A_r_*** |
| --- | --- | --- | --- | --- | --- | --- | --- | --- | --- | --- | --- |
| **Alexa fluor** | **SSR16** | F: GCACCGACGGCATTATTAGT | (AC)14 | 258-340 | 60 | 4 | 35 | 0.62 | 0.75 | 0.162 | 4.7 |
|  |  | R: GCGTAATGGCGAGCGTATT |  |  |  |  |  |  |  |  |  |
|  | **SSR19** | F: TTGACGATAAAGCGAAGGCG | (AT)13 | 184-257 | 65 | 1 | 18 | 0.56 | 0.79 | 0.296 | 4.38 |
|  |  | R: GTGGGTGCACGTACACGATA |  |  |  |  |  |  |  |  |  |
|  | **SSR2** | F: CGCCAACCGGATAATTAATTCG | (AG)19 | 187-293 | 60 | 5 | 37 | 0.91 | 0.92 | 0.004 | 6.78 |
|  |  | R: GGGAATGATTTGGAGAAGAAGTTT |  |  |  |  |  |  |  |  |  |
|  | **SSR3** | F: GTGCATATCGTATAATATCGATCGCT | (AT)18 | 228-329 | 55 | 2 | 28 | 0.86 | 0.92 | 0.065 | 6.4 |
|  |  | R: TCCTTCCAATCAGACAGTTTGA |  |  |  |  |  |  |  |  |  |
|  | **SSR49** | F: GGACCACCCTCAAACATTAAGT | (AAT)11 | 181-225 | 60 | 9 | 12 | 0.72 | 0.75 | 0.042 | 4.18 |
|  |  | R: TTTCATCAAATTGTACAGCTACACT |  |  |  |  |  |  |  |  |  |
|  | **SSR5** | F: TGCTAGCCGACATCAGATCG | (AAC)13 | 265-302 | 60 | 8 | 10 | 0.38 | 0.46 | 0.166 | 2.38 |
|  |  | R: CGTGTGACGTATTAGGTATGACC |  |  |  |  |  |  |  |  |  |
|  | **SSR6** | F: CTCCACGGACGACAACCATA | (AAT)11 | 258-300 | 60 | 7 | 18 | 0.72 | 0.75 | 0.042 | 4.54 |
|  |  | R: TAAAGACCGATGACTTGCGC |  |  |  |  |  |  |  |  |  |
|  | **SSR8** | F: CCAAGCCATCCACGCCTATG | (AC)10 | 264-287 | 60 | 6 | 12 | 0.53 | 0.54 | 0.021 | 3.26 |
|  |  | R: GCCGCGACTAAACGTTACTG |  |  |  |  |  |  |  |  |  |
| **ATTO 565** | **L4** | F: CGTGTAACTAGTATACGAACCCACC | (TGC)10  45(TGC)9 | 217-280 | 62 | 5 | 20 | 0.58 | 0.62 | 0.064 | 3.91 |
|  |  | R: GCAACAGCCATCTCCTTCTC |  |  |  |  |  |  |  |  |  |
|  | **SSR17** | F: AGTCTCCTGCTGACATTGCC | (AT)14 | 150-212 | 60 | 1 | 25 | 0.85 | 0.87 | 0.024 | 6.03 |
|  |  | R: GGTTCAAATCTAGTCGATCCGC |  |  |  |  |  |  |  |  |  |
|  | **SSR21** | F: CAACTAGGTACACGCCACGT | (AAT)12 | 176-204 | 65 | 6 | 11 | 0.67 | 0.67 | -0.002 | 3.19 |
|  |  | R: CGTTGAAATGTTCGAAATCGCG |  |  |  |  |  |  |  |  |  |
|  | **SSR38** | F: CCTGACAGCTGTGCCCTC | (AT)12 | 158-197 | 65 | 7 | 25 | 0.66 | 0.83 | 0.214 | 5.21 |
|  |  | R: GCTCGGGCGTCGTACTTATA |  |  |  |  |  |  |  |  |  |
|  | **SSR4** | F: ACAACAATTATTAATTCATCGGACCG | (AAT)17 | 164-275 | 60 | 4 | 39 | 0.84 | 0.85 | 0.02 | 5.67 |
|  |  | R: CCCAAGACAACATCGAGTCT |  |  |  |  |  |  |  |  |  |
|  | **SSR7** | F: ACGCAGGTTCCTACTGTGAT | (AT)11 | 218-256 | 60 | 2 | 11 | 0.41 | 0.59 | 0.304 | 3.33 |
|  |  | R: ACTCCTTACTGCATATACGTTTCGA |  |  |  |  |  |  |  |  |  |
|  | **SSR40** | F: TCAACCTCATTTGATGTCTTACACT | (AT)10 | 133-172 | TC - 57 | 9 | 24 | 0.72 | 0.81 | 0.12 | 5.04 |
|  |  | R: ACCATTGAGTGAACTTGGGAA |  |  |  |  |  |  |  |  |  |
| **ATTO 560** | **SSR18** | F: TTACCCATCGTGAACGCCTG | (AAT)13 | 277-305 | 65 | 5 | 9 | 0.65 | 0.65 | -0.006 | 3.5 |
|  |  | R: GGGCTTTCACTCGTGGATGT |  |  |  |  |  |  |  |  |  |
|  | **SSR23** | F: ACCGTCTTGTAAACCGCAGC | (AT)10 | 278-302 | 65 | 6 | 8 | 0.61 | 0.65 | 0.069 | 3.49 |
|  |  | R: GGCAACAGAGTTCAATCGGC |  |  |  |  |  |  |  |  |  |
|  | **SSR26** | F: CGACCGTTCAGGGAATGTTG | (AT)9 | 273-401 | 65 | 4 | 19 | 0.43 | 0.72 | 0.405 | 4.12 |
|  |  | R: AGCGCACAATCAAATATTATGGCA |  |  |  |  |  |  |  |  |  |
|  | **SSR27** | F: GAGTTATGTGCTGCATGGCG | (AAT)9 | 284-322 | 65 | 2 | 20 | 0.67 | 0.69 | 0.031 | 3.63 |
|  |  | R: CACAATCGTATTTATGTATAGGGCACT |  |  |  |  |  |  |  |  |  |
|  | **SSR28** | F: AAACCGTGACCCGAAGCG | (AC)9 | 299-344 | 65 | 1 | 10 | 0.34 | 0.65 | 0.485 | 3.16 |
|  |  | R: CCTCTTTCCCAACCAGGTGT |  |  |  |  |  |  |  |  |  |
|  | **SSR32** | F: CGTTGCCGTGTCACGTATAAT | (AC)13 | 271-383 | 65 | 7 | 45 | 0.69 | 0.79 | 0.131 | 5.42 |
|  |  | R: ATTGTCACGAGTTCGCGCTA |  |  |  |  |  |  |  |  |  |
|  | **SSR51** | F: TCATCTCTGGGACGAGGACC | (AT)12 | 279-328 | 65 | 9 | 31 | 0.32 | 0.9 | 0.646 | 5.51 |
|  |  | R: ATGTGCAGCTGAGCTCACTC |  |  |  |  |  |  |  |  |  |
| **HEX** | **SSR1** | F: CGACGCTGAGTGCCTACTAG | (AC)20 | 152-244 | 60 | 2 | 41 | 0.52 | 0.91 | 0.433 | 6.44 |
|  |  | R: GGGTCGATAGTTCAGTGTGCA |  |  |  |  |  |  |  |  |  |
|  | **SSR11** | F: ACCTGGCCATCTCACCACTA | (ACG)8 | 135-170 | 60 | 7 | 14 | 0.59 | 0.57 | -0.02 | 3.01 |
|  |  | R: CGGTTACCACCACTAAATCGGA |  |  |  |  |  |  |  |  |  |
|  | **SSR13** | F: TCGTGGTTAGTCTTAGCGAC | (AT)8 | 111-138 | 60 | 1 | 8 | 0.62 | 0.66 | 0.053 | 3.22 |
|  |  | R: ACTCACGAATAATCTCCATCACTC |  |  |  |  |  |  |  |  |  |
|  | **SSR15** | F: GGGATGAGGTGCTTCGCAA | (AC)15 | 103-208 | 60 | 5 | 50 | 0.65 | 0.74 | 0.13 | 4.66 |
|  |  | R: GTGATAGAGCGAGAATGGACGT |  |  |  |  |  |  |  |  |  |
|  | **SSR22** | F: TGTTTGGTGAAGGATCGATACG | (AG)12 | 166-208 | 65 | 4 | 16 | 0.67 | 0.72 | 0.076 | 4.15 |
|  |  | R: GGGCGATGGTTCACGGTAA |  |  |  |  |  |  |  |  |  |
|  | **SSR31** | F: ATGCAGTTGTAAATATCTATGTGGAA | (AT)14 | 143-167 | 60 | 8 | 13 | 0.82 | 0.86 | 0.044 | 5.26 |
|  |  | R: AACGGTAGTACACTAATACCTCTTT |  |  |  |  |  |  |  |  |  |
|  | **SSR46** | F: CGTGAGTTTCTAAACACCGTGG | (AC)13 | 114-159 | 65 | 9 | 16 | 0.56 | 0.83 | 0.322 | 5.08 |
|  |  | R: AGGTTGGCATGTTTGACGTAA |  |  |  |  |  |  |  |  |  |
|  | **SSR9** | F: CGGCAGATGAACCAAATCGG | (AAC)10 | 148-198 | 60 | 6 | 15 | 0.71 | 0.73 | 0.03 | 3.82 |
|  |  | R: GGCATATTATCATGTGTACACTGTAT |  |  |  |  |  |  |  |  |  |
| **-** | **Mean** | - | - | - | - | - | 21.33 | 0.63 | 0.74 | 0.15 | 4.45 |

Repeat: motifs and repeat numbers of the microsatellite markers in the individual genome from which the markers were developed; Range: allelic size range in number of base pairs (bp); T°: Annealing temperature for the PCR program; TC = Touch Down; Dye: fluorescent dyes that were used to label the forward primer; Alleles: number of alleles observed in the sample; *H_O_*: observed heterozygosity; *H_E_*: expected heterozygosity; *F_IS_*: fixation index; *A_R_*: allelic richness.

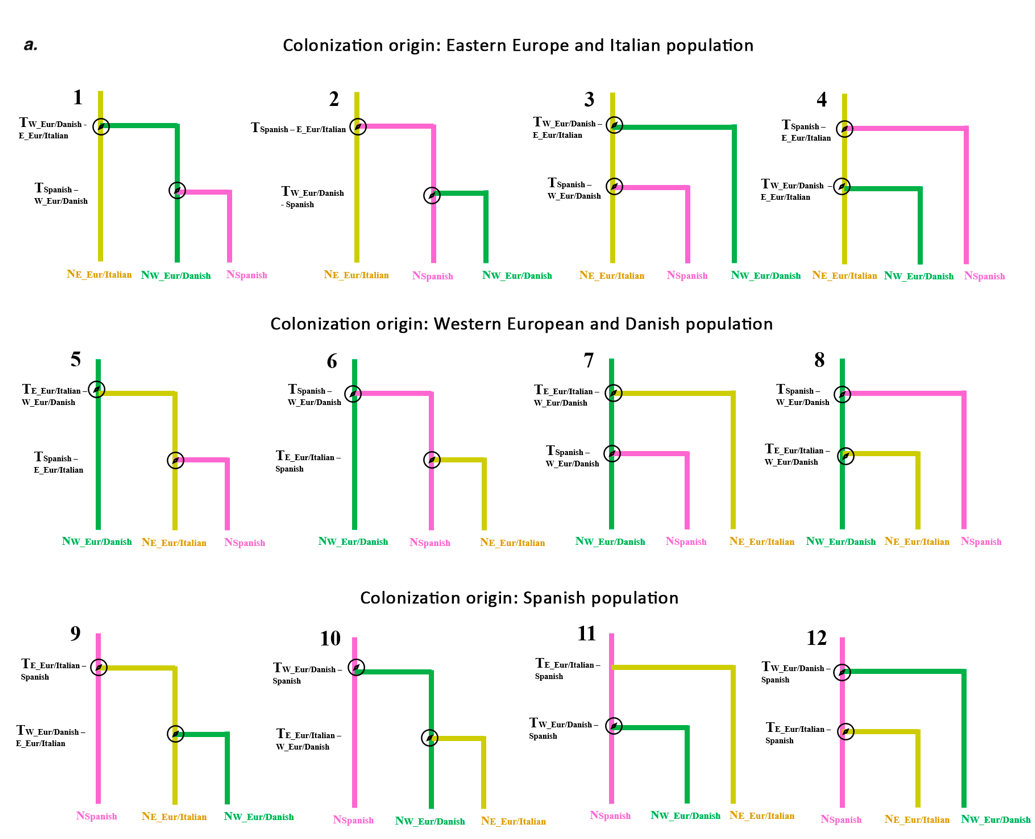

**Figure S2.** **Demographic models compared with the random-forest approximate Bayesian computation (ABC-RF) to infer the colonization history of the rosy apple aphid (29 perfect-motif microsatellite markers).** a**. The first ABC-RF step inferred the colonization history in Europe and included the European populations detected with STRUCTURE for *K*=5: the Spanish (pink, *N*=113 individuals), the Eastern European and the Italian (“E_Eur/Italian”; yellow, *N*=114), the Western European and Danish ((“W_Eur/Danish”; green, *N*=89). For step 1, each model was run with gene flow among populations and without gene flow; thus 24 models were run.** **b.** The second ABC-RF step inferred the colonization of the rosy apple aphid out of Europe, and included the European population (merging the Spanish, Eastern European and Italian, Western European and Danish populations, *N*=316), the Moroccan (blue, *N*=63), and the US (orange, *N*=28). For step 2, each model was run with five gene flow modalities: no gene flow, gene flow among all populations, gene flow between each pair of population; therefore 30 models were run. *T_X-Y_*: divergence time between *Y* and *X* population; *N_X_*: effective population size.

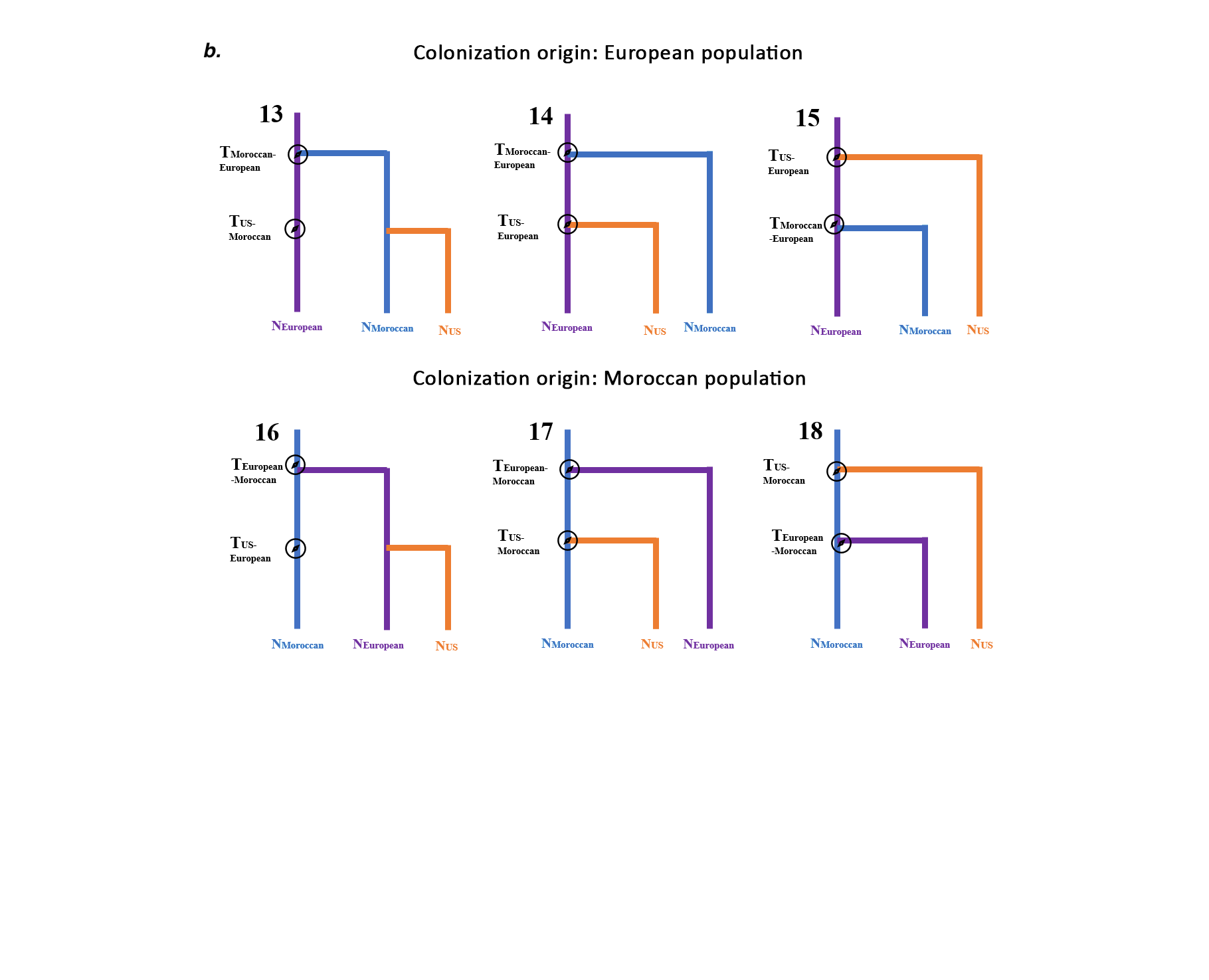

**Figure S2 (continued).** **Demographic models compared with the random-forest approximate Bayesian computation (ABC-RF) to infer the colonization history of the rosy apple aphid (29 perfect-motif microsatellite markers).** a. The first ABC-RF step inferred the colonization history in Europe and included the European populations detected with STRUCTURE for *K*=5: the Spanish (pink, *N*=113 individuals), the Eastern European and Italian (“E_Eur/Italian”; yellow, *N*=114); the Western European and Danish ((“W_Eur/Danish”; green, *N*=89). For step 1, each model was run with gene flow among populations and without gene flow; thus 24 models were run. **b. The second ABC-RF step inferred the colonization of the rosy apple aphid out of Europe, and included the European population (merging the Spanish, Eastern European and Italian, Western European and Danish populations, *N*=316), the Moroccan (blue, *N*=63), and the US (orange, *N*=28). For step 2, each model was run with five gene flow modalities: no gene flow, gene flow among all populations, gene flow between each pair of population; therefore 30 models were run. *T_X-Y_*: divergence time between *Y* and *X* population; *N_X_*: effective population size.**

**Table S3.** **Prior distributions for each model parameter used for Random Forest approximate Bayesian computations to reconstruct *Dysaphis plantaginea* colonization history.** Prior distributions are uniform or log-uniform between lower and upper bounds.

| **Parameter** | **Distribution** | **Lower bound** | **Upper bound** |
| --- | --- | --- | --- |
| *N_E_* | *unif* | 50 | 100,000 |
| *Tx-y* (in years) | *logunif* | 1 | 10,000 |
| *μ* | *logunif* | 1.00E-07 | 1.00E-04 |
| *ARG* | *unif* | 1 | 30 |
| *Growth rate* | *unif* | -0.5 | 0 |

*N_E_*: effective population size, *T_X-Y_*: divergence time between the *X* and *Y* populations, *μ*: mutation rate, unif: uniform distribution, *log-unif*: log-uniform distribution.

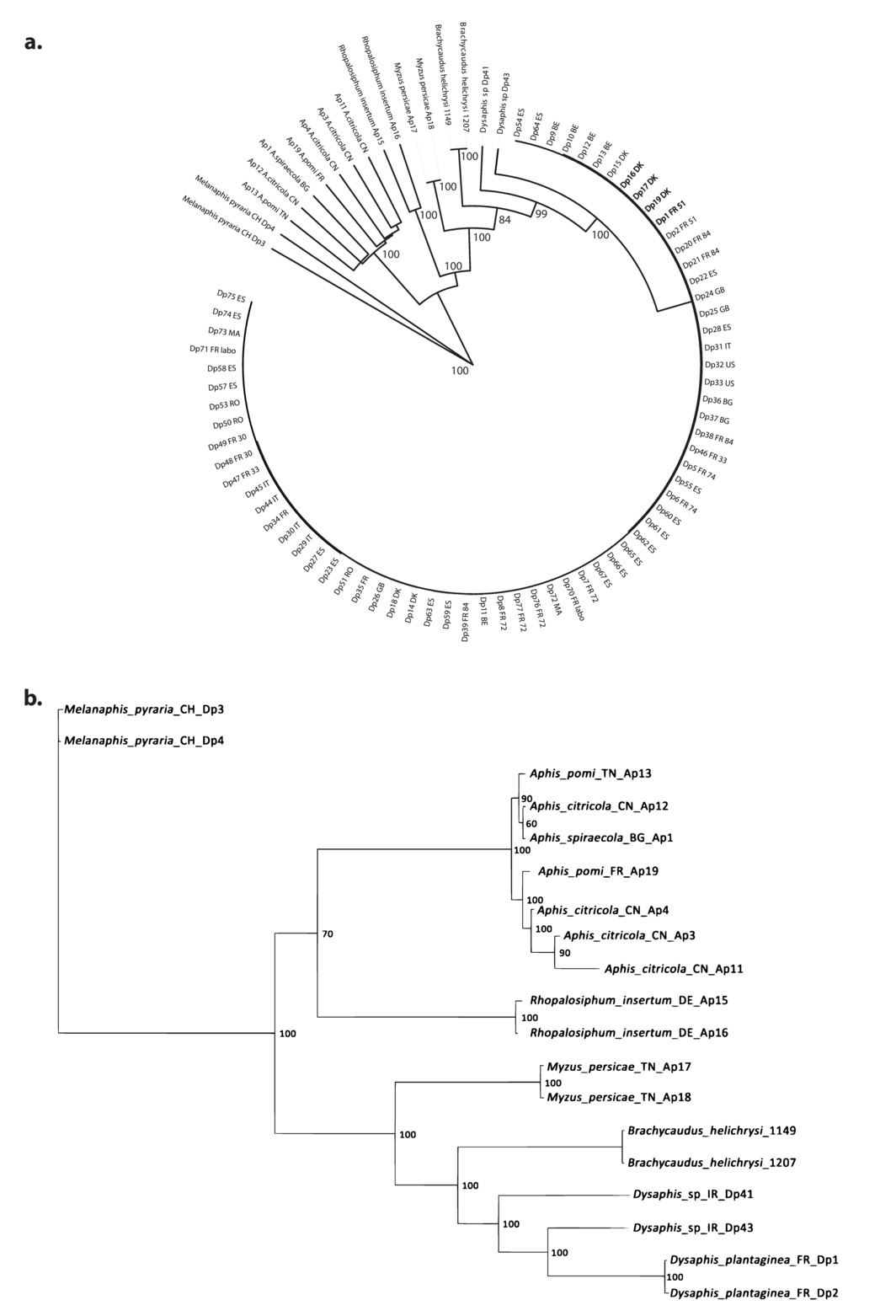

**Figure S3. Phylogenetic relationships between *Dysaphis plantaginea* and other aphid species found on fruit trees inferred with MrBayes using the concatenated sequences of CO1, CytB, and TrpB markers a.** including the 86 individuals belonging to the *Dysaphis*, *Aphis*, *Melanaphis*, *Rhopalosiphum,* and *Myzus* genera. **b.** including only two representants of *D. plantaginea* and other species**.** Note that two sequences of *Brachycaudus* *helisychri* from (Popkin et al., 2017) were added. Bootstrap values were obtained after 1,000 repetitions, and only bootstrap > 80% are shown. Each sample is represented with the species ID: DP for *D. plantaginea*, ID number, and a 2-digit country abbreviation (BE: Belgium, CH: Switzerland, CN: China, DK: Denmark, ES: Spain, FR: France, GB: UK, IT: Italy, MA: Morocco, RO: Romania, TN: Tunisia).

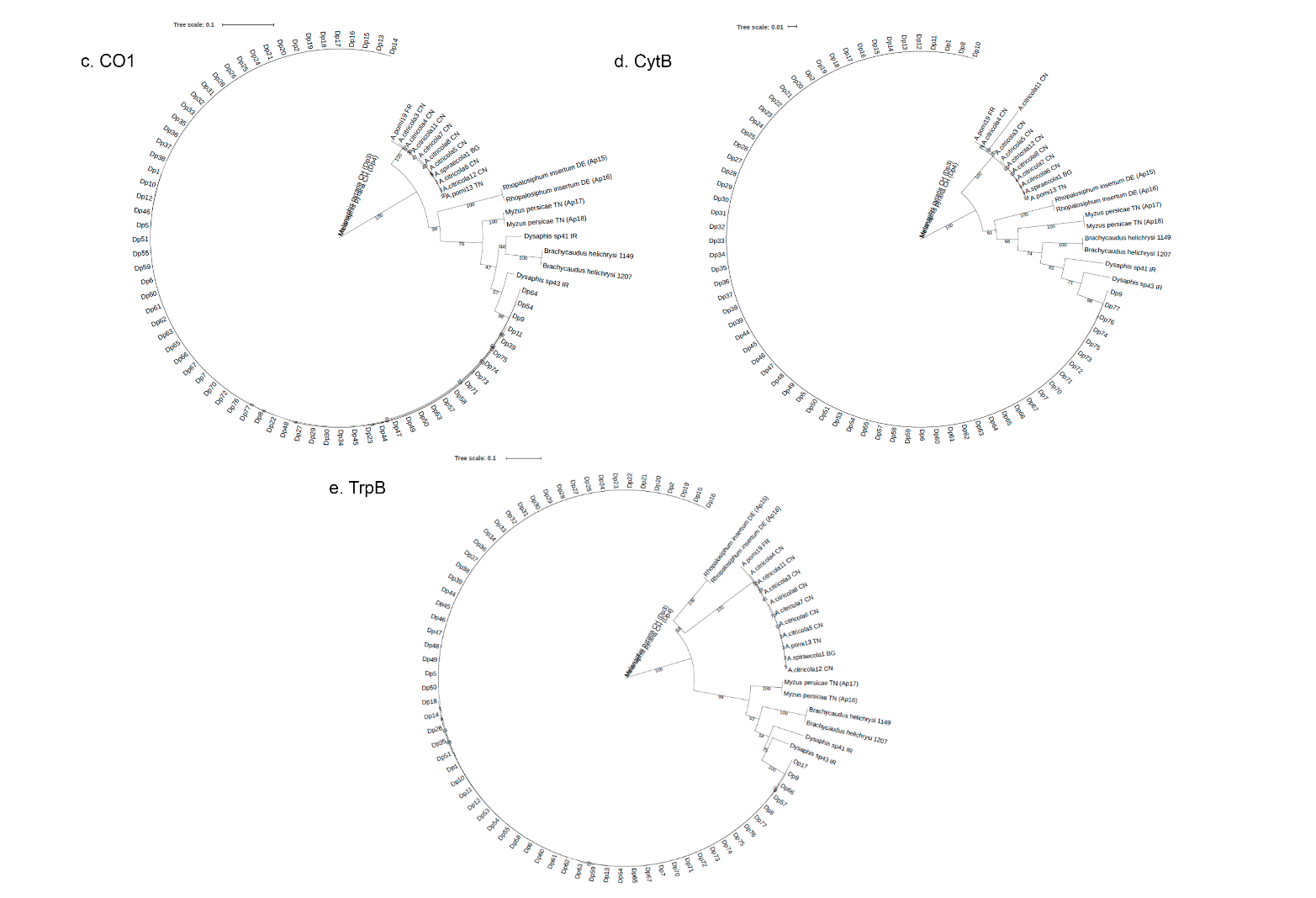

**Figure S3 (continued). Phylogenetic relationships between *Dysaphis plantaginea* and other aphid species found on fruit trees inferred with RAxML using three different markers.** Phylogenetic trees inferred from c. CO1, d. CytB, and e. TrpB markers. The sequences were extracted from 86 individuals of *Dysaphi*s (Dp), *Aphis*, *Melanaphis*, *Rhopalosiphum*, *Myzus* genera, and two sequences of *Brachycaudus helisychri* from (Popkin et al., 2017). Bootstrap values were obtained after 1,000 repetitions, and only bootstrap > 80% are shown. Each sample is represented with the species ID: DP for *Dysaphis plantaginea*, ID number, and a 2-digit country abbreviation (BE: Belgium, CH: Switzerland, CN: China, DK: Denmark, ES: Spain, FR: France, GB: the UK, IT: Italy, MA: Morocco, RO: Romania, TN: Tunisia).

| **Site** | **Site** | **Country** | **Host** | **Apple variety** | ***N*** | ***G*** | ***G/N*** | ***H_o_*** | ***H_e_*** | ***F_is_*** | ***P-value*** | ***Ar*** | ***Ap*** | **Y** | **X** |
| --- | --- | --- | --- | --- | --- | --- | --- | --- | --- | --- | --- | --- | --- | --- | --- |
| **GIN_1** | Gingelom | Belgium | *Malus domestica* | Topaz | 13 | 6 | 0.46 | 0.63 | 0.64 | 0.01 | 0.29 | 4.19 | 0.13 | 50.762 | 5.153 |
| **GIN_2** |  |  |  | Golden | 15 | 5 | 0.33 | 0.63 | 0.59 | -0.07 | 1 | 3.53 | 0.1 | 50.763 | 5.155 |
| **GIN_3** |  |  |  | Jonagold king | 15 | 3 | 0.2 | 0.75 | 0.38 | -0.97 | 1 | - | - | 50.763 | 5.154 |
| **GIN_4** |  |  |  | Topaz | 15 | 10 | 0.67 | 0.56 | 0.62 | 0.11 | 0 | 4.08 | 0.05 | 50.762 | 5.154 |
| **STT_1** | Sint-Truiden |  |  | Novajo | 13 | 7 | 0.54 | 0.65 | 0.65 | 0.01 | 0.16 | 4.24 | 0.12 | 50.772 | 5.157 |
| **STT_2** |  |  |  | Wellant | 14 | 1 | 0.07 | 0.52 | 0.26 | -1 | 1 | - | - | 50.764 | 5.154 |
| **STT_3** |  |  |  | Boscoop | 14 | 3 | 0.21 | 0.64 | 0.44 | -0.48 | 1 | - | - | 50.773 | 5.157 |
| **STT_4** |  |  |  | Jonagored | 14 | 5 | 0.36 | 0.76 | 0.63 | -0.22 | 1 | 3.17 | 0.16 | 50.766 | 5.155 |
| **PLO** | Plovdiv | Bulgari |  | NA | 15 | 10 | 0.67 | 0.63 | 0.7 | 0.1 | 0 | 4.62 | 0.09 | 42.15 | 24.75 |
| **DAR** | Darmstadt | Germany |  | NA | 7 | 7 | 1 | 0.63 | 0.68 | 0.07 | 0.11 | 4.41 | 0.19 | 49.873 | 8.651 |
| **FAX** | Faxe | Denmark |  | Holsteiner Cox & Red Ingrid Marie | 13 | 13 | 1 | 0.63 | 0.76 | 0.16 | 0 | 4.91 | 0.18 | 55.187 | 12.112 |
| **LIL** | Lille Skensved |  |  | Aroma | 15 | 15 | 1 | 0.69 | 0.76 | 0.09 | 0 | 5.09 | 0.12 | 55.52 | 12.136 |
| **TAA** | Taastrup |  |  | Fillipa & Holsteiner Cox & Grasten Yellow & Grasten Red | 14 | 14 | 1 | 0.67 | 0.78 | 0.13 | 0 | 4.95 | 0.15 | 55.673 | 12.308 |
| **CAM** | Camoca | Spain |  | NA | 11 | 10 | 0.91 | 0.61 | 0.68 | 0.11 | 0 | 4.37 | 0.1 | 43.453 | -5.483 |
| **LAS** | La Salve |  |  | NA | 15 | 15 | 1 | 0.65 | 0.73 | 0.1 | 0 | 4.55 | 0.06 | 43.381 | -5.65 |
| **LLE** | Lleida |  |  | Golden 972 | 14 | 14 | 1 | 0.65 | 0.73 | 0.11 | 0 | 4.38 | 0.11 | 41.587 | 0.583 |
| **PRI** | Priesca |  |  | NA | 15 | 15 | 1 | 0.55 | 0.68 | 0.18 | 0 | 4.3 | 0.03 | 43.485 | -5.361 |
| **SEL** | Selorio |  |  | NA | 12 | 12 | 1 | 0.58 | 0.68 | 0.15 | 0 | 4.34 | 0.08 | 43.516 | -5.348 |
| **SIE** | Siero |  |  | NA | 9 | 9 | 1 | 0.62 | 0.68 | 0.09 | 0 | 4.35 | 0.06 | 43.389 | -5.584 |
| **SOR** | Sorribes |  |  | NA | 15 | 15 | 1 | 0.62 | 0.7 | 0.1 | 0 | 4.46 | 0.07 | 43.481 | -5.446 |
| **VIL_1** | Villaviciosa |  |  | NA | 13 | 13 | 1 | 0.6 | 0.7 | 0.15 | 0 | 4.45 | 0.1 | 43.478 | -5.442 |
| **VIL_2** | Poisy |  |  | NA | 15 | 15 | 1 | 0.6 | 0.69 | 0.12 | 0 | 4.41 | 0.08 | 43.474 | -5.444 |
| **AVI_1** | Avignon | France |  | Grany | 10 | 10 | 1 | 0.58 | 0.7 | 0.17 | 0 | 4.44 | 0.03 | 43.917 | 4.881 |
| **AVI_2** | Cobham |  |  | Ariane | 11 | 11 | 1 | 0.69 | 0.73 | 0.05 | 0.01 | 4.76 | 0.07 | 43.914 | 4.882 |
| **BEL** | Bellegarde |  |  | Ariane | 14 | 14 | 1 | 0.62 | 0.74 | 0.16 | 0 | 4.89 | 0.11 | 43.74 | 4.45 |
| **BON_1** | Bonnetable |  |  | Belle de Boskoop & Elista & Melrose & Jonagold | 12 | 12 | 1 | 0.64 | 0.76 | 0.15 | 0 | 4.88 | 0.11 | 48.179 | 0.425 |
| **BON_2** | Asni |  |  | NA | 14 | 14 | 1 | 0.66 | 0.74 | 0.11 | 0 | 4.65 | 0.1 | 48.179 | 0.425 |
| **BON_3** | Ait Ayache |  |  | NA | 14 | 14 | 1 | 0.63 | 0.75 | 0.16 | 0 | 4.78 | 0.16 | 48.179 | 0.425 |
| **BON_4** | Ait Bouguemez |  |  | NA | 15 | 15 | 1 | 0.64 | 0.76 | 0.16 | 0 | 4.91 | 0.1 | 48.179 | 0.425 |
| **GOT_1** | Gotheron |  |  | Eco-Ariane & Eco-Melrose & Bio-Smoothie & Eco-Smoothie | 15 | 15 | 1 | 0.64 | 0.73 | 0.11 | 0 | 4.58 | 0.06 | 44.977 | 4.93 |
| **GOT_2** | Naour |  |  | NA | 10 | 10 | 1 | 0.63 | 0.75 | 0.16 | 0 | 4.57 | 0.07 | 44.977 | 4.93 |
| **GOT_3** | Sighisoara |  |  | NA | 12 | 12 | 1 | 0.65 | 0.74 | 0.12 | 0 | 4.78 | 0.09 | 44.977 | 4.93 |
| **GOT_4** | Tauti |  |  | NA | 15 | 15 | 1 | 0.64 | 0.72 | 0.12 | 0 | 4.66 | 0.1 | 44.977 | 4.93 |
| **LOO** | Loos-en-Gohelle |  |  | Jonagored | 15 | 14 | 0.93 | 0.64 | 0.73 | 0.12 | 0 | 4.84 | 0.15 | 50.458 | 2.793 |
| **NOI** | Noirlieu |  |  | Boskoop & Jonagold & Reinette | 13 | 11 | 0.85 | 0.69 | 0.73 | 0.05 | 0 | 4.79 | 0.11 | 48.95 | 4.81 |
| **POI** | Poisy |  |  | Idared | 7 | 7 | 1 | 0.65 | 0.72 | 0.09 | 0.01 | 4.43 | 0.06 | 45.921 | 6.064 |
| **TOU** | Toulenne |  |  | Val & Akane & Choupette & Topaze | 13 | 13 | 1 | 0.63 | 0.74 | 0.14 | 0 | 4.65 | 0.08 | 44.559 | -0.262 |
| **COB** | Cobham | UK |  | 17 different varieties | 16 | 16 | 1 | 0.69 | 0.77 | 0.1 | 0 | 4.81 | 0.08 | 51.399 | 0.392 |

**Table S7.** **Genetic diversity estimates, geographic origin, and host species for the 52 *Dysaphis plantaginea* sampling sites (*N*=667, 30 microsatellite markers).**

**Table S7 (continued).** **Genetic diversity estimates, geographic origin, and host species for the 52 *Dysaphis plantaginea* sampling sites (*N*=667, 30 microsatellite markers).**

| **Site** | **Site** | **Country** | **Host** | **Apple variety** | ***N*** | ***G*** | ***G/N*** | ***H_o_*** | ***H_e_*** | ***F_IS_*** | ***P-value*** | ***Ar*** | ***Ap*** | **Y** | **X** |
| --- | --- | --- | --- | --- | --- | --- | --- | --- | --- | --- | --- | --- | --- | --- | --- |
| **CAD** | Cadriano | Italy | *Malus domestica* | Reinette de Champagne | 13 | 13 | 1 | 0.67 | 0.75 | 0.12 | 0 | 4.75 | 0.08 | 44.51 | 11.41 |
| **FER** | Ferrara |  |  | Pink Lady | 12 | 12 | 1 | 0.66 | 0.75 | 0.12 | 0 | 4.92 | 0.38 | 44.87 | 11.49 |
| **ASN** | Asni | Morocco |  | Gala Brookfield & Golden & Buckeye | 13 | 13 | 1 | 0.51 | 0.55 | 0.08 | 0 | 3.47 | 0.04 | 31.242 | -7.987 |
| **AYA** | Ait Ayache |  |  | Gala & Golden Delicious & Starking Delicious | 15 | 15 | 1 | 0.51 | 0.61 | 0.15 | 0 | 3.73 | 0.03 | 32.667 | -4.935 |
| **BOU** | Ait Bouguemez |  |  | NA | 15 | 15 | 1 | 0.54 | 0.57 | 0.06 | 0 | 3.63 | 0.02 | 31.646 | -6.468 |
| **IMO** | Imouzzer Kandar |  |  | Gala & Golden Reinders & Golden Smoothee | 9 | 9 | 1 | 0.58 | 0.65 | 0.09 | 0 | 3.9 | 0.1 | 33.731 | -5.016 |
| **NAO** | Naour |  |  | NA | 11 | 11 | 1 | 0.56 | 0.62 | 0.09 | 0 | 3.8 | 0.07 | 32.48 | -5.948 |
| **SIG** | Sighisoara | Romania |  | NA | 14 | 14 | 1 | 0.66 | 0.75 | 0.12 | 0 | 4.96 | 0.15 | 46.22 | 24.796 |
| **TAU** | Tauti |  |  | Goldspur & Gloster & Aromat de Vara & Mutsu & William's Pride | 10 | 10 | 1 | 0.66 | 0.74 | 0.11 | 0 | 4.77 | 0.11 | 46.712 | 23.506 |
| **GEN** | Geneva | USA |  | Red Delicious & Cortland & Empire & Jonagold | 15 | 14 | 0.93 | 0.57 | 0.64 | 0.11 | 0 | 3.74 | 0.08 | 42.866 | -77.025 |
| **WIN** | Winchester |  |  | Smoothie Golden | 15 | 15 | 1 | 0.58 | 0.63 | 0.09 | 0 | 3.7 | 0.04 | 39.112 | -78.287 |
| **LOO_P** | Loos-en-Gohelle | France | *Plantago lanceolata* | NA | 6 | 5 | 0.83 | 0.73 | 0.74 | 0.02 | 0.27 | 4.67 | 0.17 | 50.458 | 2.793 |
| **LAB** | Reared in lab |  |  | NA | 10 | 9 | 0.9 | 0.66 | 0.7 | 0.05 | 0 | 4.52 | 0.13 | - | - |
| **ALT** | Alta Ribagorça | Spain | *Malus domestica* | NA | 7 | 7 | 1 | 0.65 | 0.71 | 0.09 | 0 | 4.5 | 0.09 | 42.448 | 0.707 |
| **TOT** |  |  |  |  | 667 | 582 | 0.87 | 0.6273 | 0.7386 | 0.1506 | 0 | 4.4346 | 0.1004 |  |  |

Site: orchard of aphid population sampling; Country: country of aphid population origin; N: number of individuals; G: number of multilocus genotypes (*MLG*); G/N: proportion of unique MLG; *H_O_:* observed heterozygosity; *H_E_*: expected heterozygosity; *F_IS_*: Fixation index; P-values: P-values associated with the *F_IS_;* *Ar:* allelic richness; *Ap*: private allelic richness; Y and X: latitude and longitude of the sampling site. Bold value is significant.

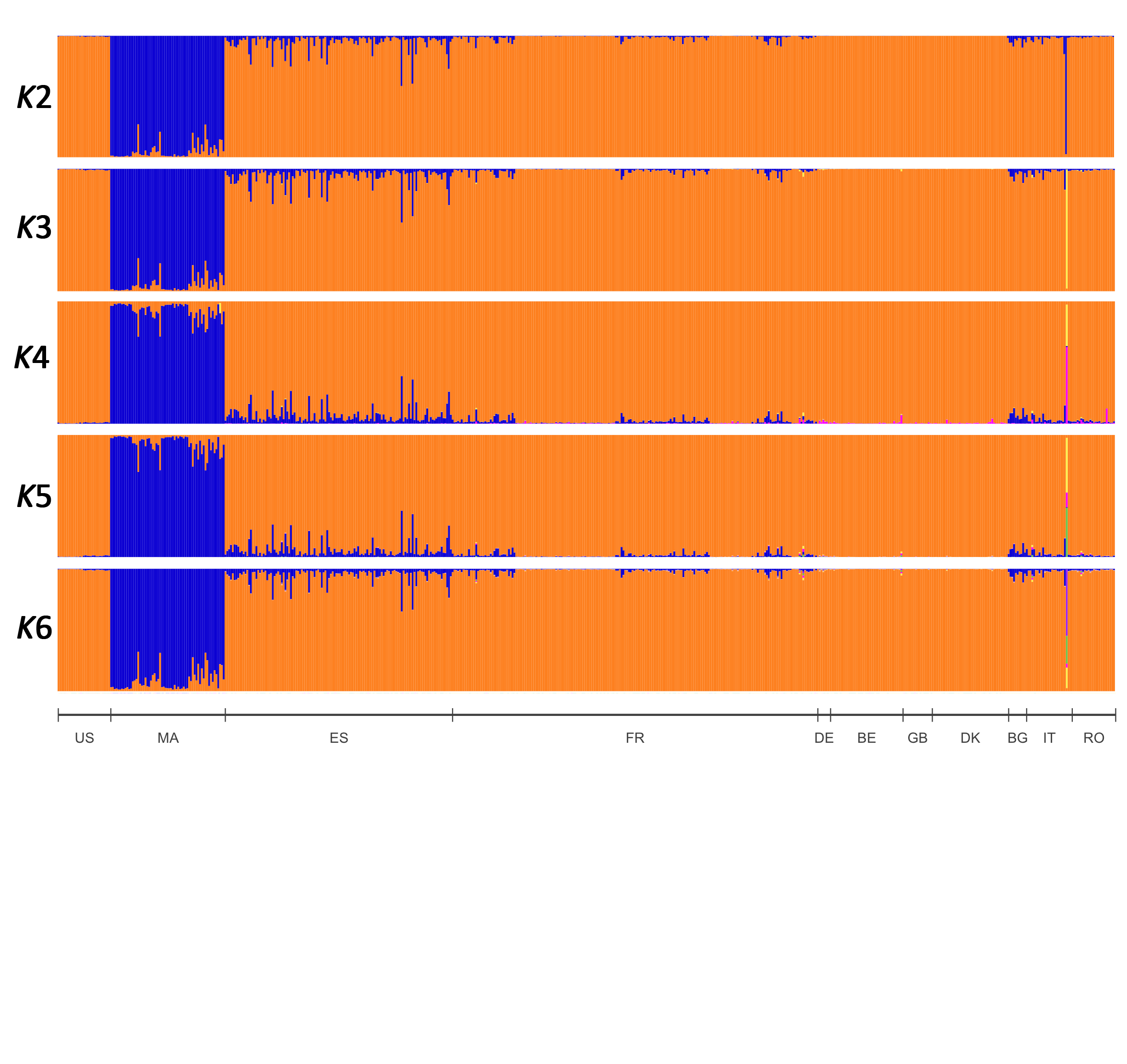

**Figure S4. Spatial population genetic structure of the rosy apple aphid *Dysaphis plantaginea* in Europe, the Mediterranean Basin, and the US (*N* = 582 individuals, 52 sites), inferred with TESS from *K* = 2 to *K* = 6 (for K>6 did not reveal additional structure).** Each individual is represented by a vertical bar, partitioned into *K* segments representing the proportions of the ancestry of its genome in *K* clusters. For a better visualization, the 52 sites were grouped by country and ordered according to a latitudinal gradient (except for the US which is positioned before the others). US: United States of America; MA: Morocco; ES: Spain; BG: Bulgaria; IT: Italy; FR: France; RO: Romania; DE: Germany; BE: Belgium; GB: United Kingdom; DK: Denmark.

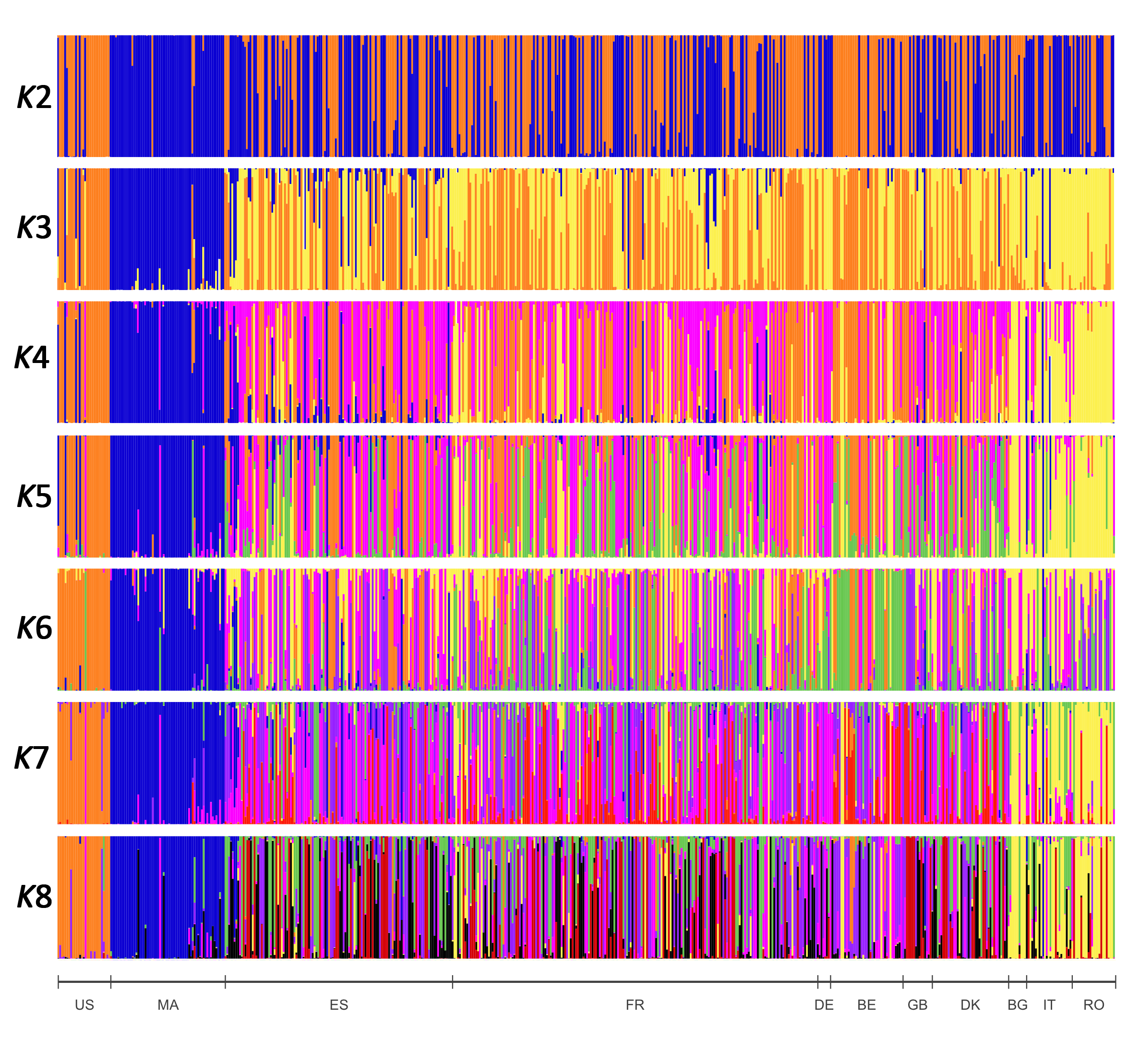

**Figure S5.** **Population structure of the rosy apple aphid *Dysaphis plantaginea* in Europe, the Mediterranean Basin, and the US (*N* = 582 individuals, 52 sites), inferred with the Discriminant Analysis of Principal Components from *K* = 2 to *K* = 8.** Each individual is represented by a vertical bar, partitioned into *K* segments representing the proportions of the ancestry of its genome in *K* clusters. For better visualization, the 52 sites were grouped by country and ordered according to a latitudinal gradient (except for the US which is positioned before the others). US: United States of America; MA: Morocco; BG: Bulgaria; IT: Italy; RO: Romania; ES: Spain; FR: France; DE: Germany; BE: Belgium; GB: United Kingdom; DK: Denmark.

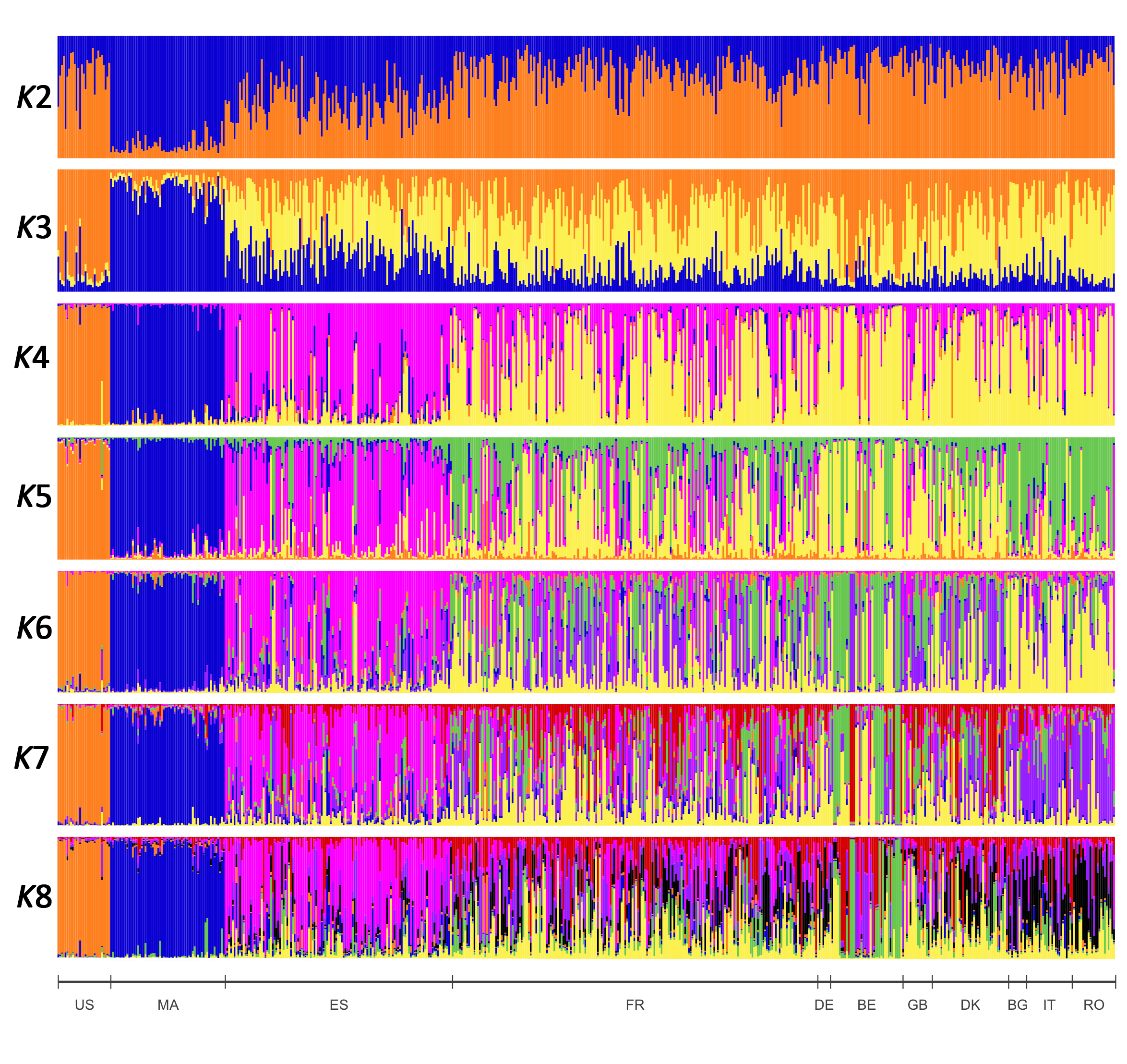

**Figure S6.** **Spatial population genetic structure of the rosy apple aphid *Dysaphis plantaginea* in Europe, the Mediterranean Basin, and the US (*N* = 582 individuals, 52 sites), inferred with STRUCTURE from *K* = 2 to *K* = 8.** Each individual is represented by a vertical bar, partitioned into *K* segments representing the proportions of the ancestry of its genome in *K* clusters. When several clustering solutions (‘modes’) were found within replicate runs, we only show the major mode. For better visualization, the 52 sites were grouped by country and ordered according to a latitudinal gradient (except for the US which is positioned before the others). US: United States of America; MA: Morocco; ES: Spain; BG: Bulgaria; IT: Italy; FR: France; RO: Romania; DE: Germany; BE: Belgium; GB: United Kingdom; DK: Denmark.

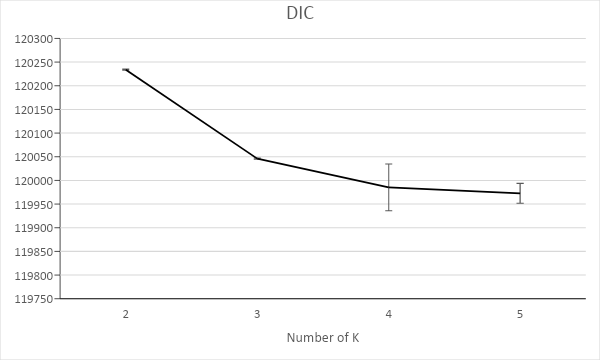

**Figure S7.** Estimated number of populations for *Dysaphis plantaginea* from TESS results using the Discriminant Information Criterion (*DIC*).

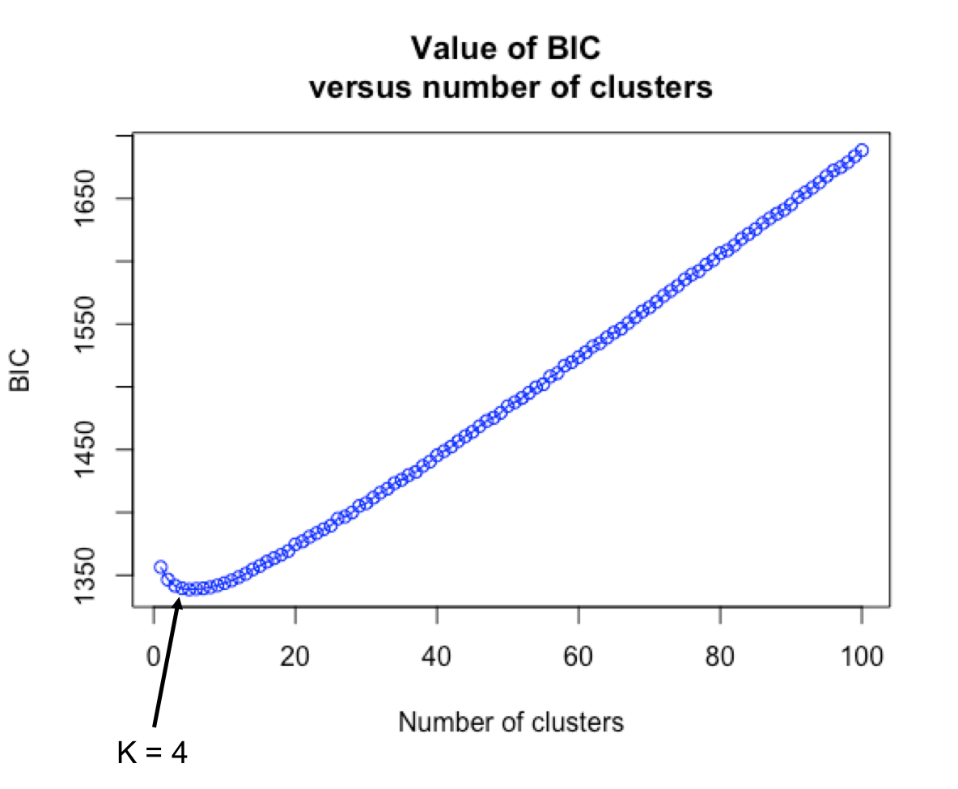

**Figure S8.** Estimated number of populations in *Dysaphis plantaginea* with DAPC analyses using the Bayesian Information Criterion (*BIC*).

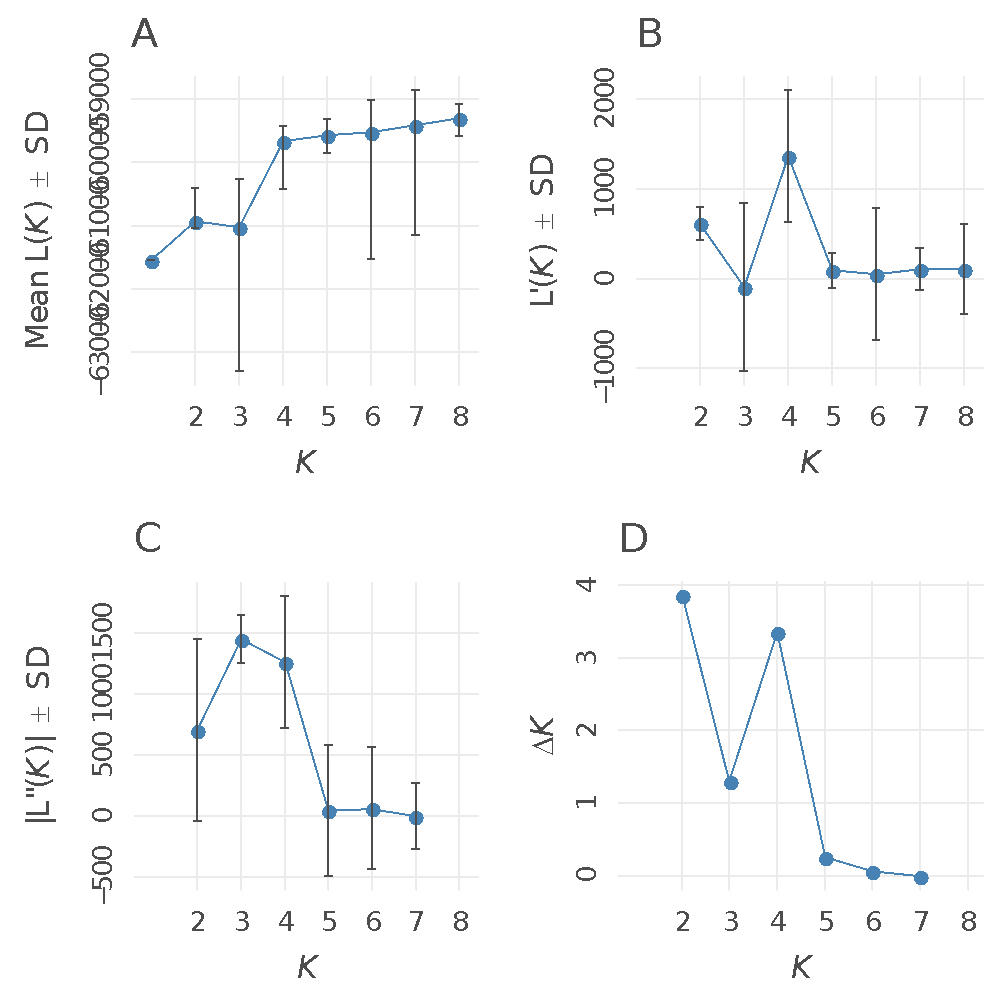

**Figure S9.** Estimated number of populations for *Dysaphis plantaginea* estimated with STRUCTURE using the plot of *∆K* in function of the assumed *K* value.

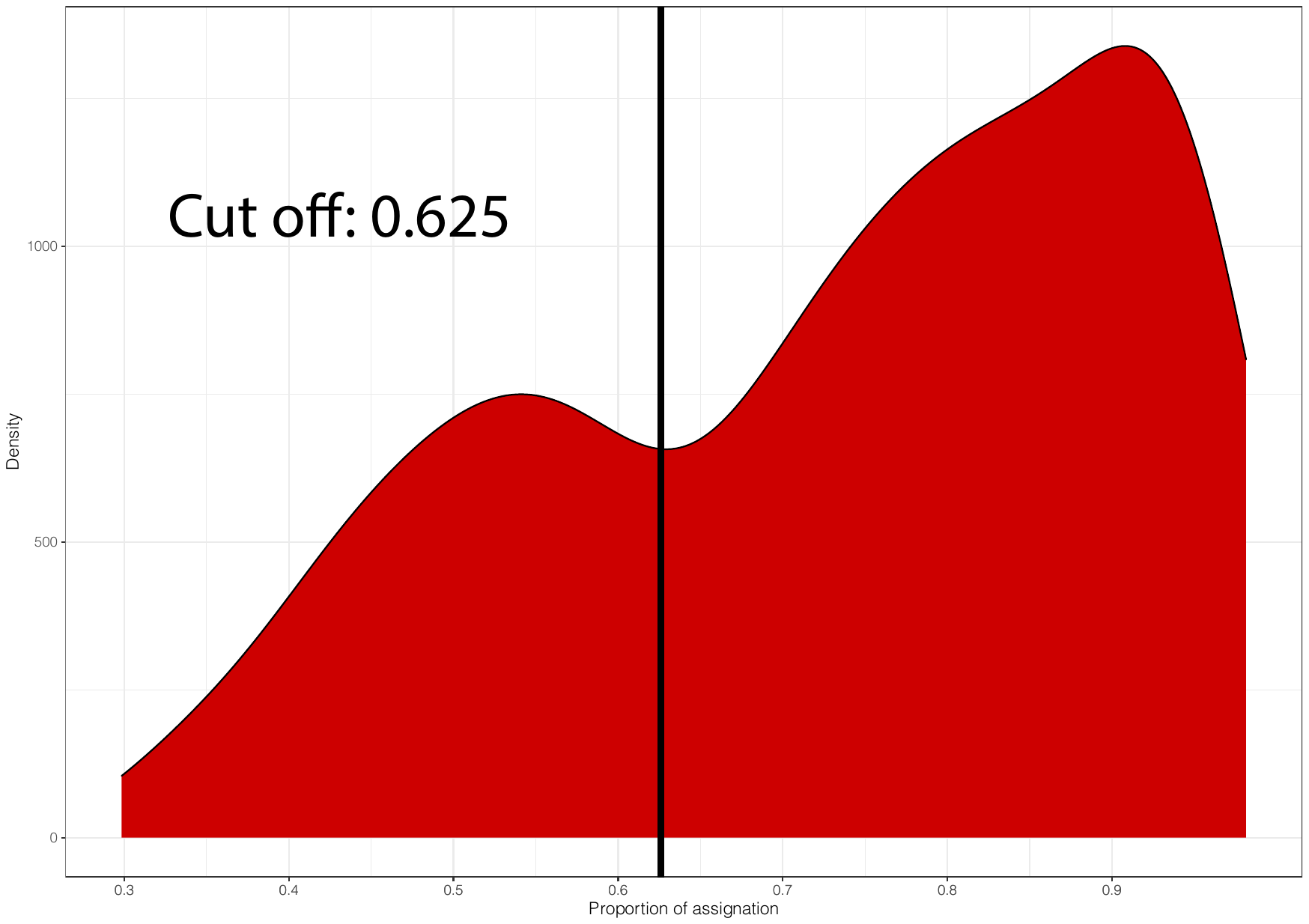

**Figure S10.** Distribution of the maximum membership coefficient inferred with STRUCTURE for the 582 individuals of *Dysaphis plantaginea* for *K*=5. The vertical line at 0.625 represents the threshold used to assign an individual to a given cluster. Individuals with a membership coefficient < 0.625 were considered as admixed genotypes. The y-axis represents the number of individuals while the x-axis presents the maximum membership coefficient of each individual to a given genetic group inferred with STRUCTURE.

**Table S8. Genetic diversity statistics, geographic origin, and host species of 52 sampling sites of *Dysaphis plantaginea* colonies**

| **Country** | **Site** | **Site** | **Host** | **Apple variety** | ***N*** | ***G*** | ***G/N*** | ***H_o_*** | ***H_e_*** | ***F_is_*** | ***P-value*** | ***Ar*** | ***Ap*** | **Y** | **X** |
| --- | --- | --- | --- | --- | --- | --- | --- | --- | --- | --- | --- | --- | --- | --- | --- |
| **Belgium** | **GIN_1** | Gingelom | *Malus domestica* | Topaz | 13 | 6 | 0.46 | 0.63 | 0.64 | 0.01 | 0.29 | 4.19 | 0.13 | 50.762 | 5.153 |
|  | **GIN_2** |  |  | Golden | 15 | 5 | 0.33 | 0.63 | 0.59 | -0.07 | 1 | 3.53 | 0.1 | 50.763 | 5.155 |
|  | **GIN_3** |  |  | Jonagold king | 15 | 3 | 0.2 | 0.75 | 0.38 | -0.97 | 1 | - | - | 50.763 | 5.154 |
|  | **GIN_4** |  |  | Topaz | 15 | 10 | 0.67 | 0.56 | 0.62 | 0.11 | 0 | 4.08 | 0.05 | 50.762 | 5.154 |
|  | **STT_1** | Sint-Truiden |  | Novajo | 13 | 7 | 0.54 | 0.65 | 0.65 | 0.01 | 0.16 | 4.24 | 0.12 | 50.772 | 5.157 |
|  | **STT_2** |  |  | Wellant | 14 | 1 | 0.07 | 0.52 | 0.26 | -1 | 1 | - | - | 50.764 | 5.154 |
|  | **STT_3** |  |  | Boscoop | 14 | 3 | 0.21 | 0.64 | 0.44 | -0.48 | 1 | - | - | 50.773 | 5.157 |
|  | **STT_4** |  |  | Jonagored | 14 | 5 | 0.36 | 0.76 | 0.63 | -0.22 | 1 | 3.17 | 0.16 | 50.766 | 5.155 |
| **Bulgaria** | **PLO** | Plovdiv |  | NA | 15 | 10 | 0.67 | 0.63 | 0.7 | 0.1 | 0 | 4.62 | 0.09 | 42.15 | 24.75 |
| **Germany** | **DAR** | Darmstadt |  | NA | 7 | 7 | 1 | 0.63 | 0.68 | 0.07 | 0.11 | 4.41 | 0.19 | 49.873 | 8.651 |
| **Denmark** | **FAX** | Faxe |  | Holsteiner Cox & Red Ingrid Marie | 13 | 13 | 1 | 0.63 | 0.76 | 0.16 | 0 | 4.91 | 0.18 | 55.187 | 12.112 |
|  | **LIL** | Lille Skensved |  | Aroma | 15 | 15 | 1 | 0.69 | 0.76 | 0.09 | 0 | 5.09 | 0.12 | 55.52 | 12.136 |
|  | **TAA** | Taastrup |  | Fillipa & Holsteiner Cox & Grasten Yellow & Grasten Red | 14 | 14 | 1 | 0.67 | 0.78 | 0.13 | 0 | 4.95 | 0.15 | 55.673 | 12.308 |
| **Spain** | **CAM** | Camoca |  | NA | 11 | 10 | 0.91 | 0.61 | 0.68 | 0.11 | 0 | 4.37 | 0.1 | 43.453 | -5.483 |
|  | **LAS** | La Salve |  | NA | 15 | 15 | 1 | 0.65 | 0.73 | 0.1 | 0 | 4.55 | 0.06 | 43.381 | -5.65 |
|  | **LLE** | Lleida |  | Golden 972 | 14 | 14 | 1 | 0.65 | 0.73 | 0.11 | 0 | 4.38 | 0.11 | 41.587 | 0.583 |
|  | **PRI** | Priesca |  | Local cider cultivars | 15 | 15 | 1 | 0.55 | 0.68 | 0.18 | 0 | 4.3 | 0.03 | 43.485 | -5.361 |
|  | **SEL** | Selorio |  | Local cider cultivars | 12 | 12 | 1 | 0.58 | 0.68 | 0.15 | 0 | 4.34 | 0.08 | 43.516 | -5.348 |
|  | **SIE** | Siero |  | Local cider cultivars | 9 | 9 | 1 | 0.62 | 0.68 | 0.09 | 0 | 4.35 | 0.06 | 43.389 | -5.584 |
|  | **SOR** | Sorribes |  | Local cider cultivars | 15 | 15 | 1 | 0.62 | 0.7 | 0.1 | 0 | 4.46 | 0.07 | 43.481 | -5.446 |
|  | **VIL_1** | Villaviciosa |  | Local cider cultivars | 13 | 13 | 1 | 0.6 | 0.7 | 0.15 | 0 | 4.45 | 0.1 | 43.478 | -5.442 |
|  | **VIL_2** | Villaviciosa |  | Local cider cultivars | 15 | 15 | 1 | 0.6 | 0.69 | 0.12 | 0 | 4.41 | 0.08 | 43.474 | -5.444 |
| **France** | **AVI_1** | Avignon |  | Grany | 10 | 10 | 1 | 0.58 | 0.7 | 0.17 | 0 | 4.44 | 0.03 | 43.917 | 4.881 |
|  | **AVI_2** | Cobham |  | Ariane | 11 | 11 | 1 | 0.69 | 0.73 | 0.05 | 0.01 | 4.76 | 0.07 | 43.914 | 4.882 |
|  | **BEL** | Bellegarde |  | Ariane | 14 | 14 | 1 | 0.62 | 0.74 | 0.16 | 0 | 4.89 | 0.11 | 43.74 | 4.45 |
|  | **BON_1** | Bonnetable |  | Belle de Boskoop & Elista & Melrose & Jonagold | 12 | 12 | 1 | 0.64 | 0.76 | 0.15 | 0 | 4.88 | 0.11 | 48.179 | 0.425 |
|  | **BON_2** | Asni |  |  | 14 | 14 | 1 | 0.66 | 0.74 | 0.11 | 0 | 4.65 | 0.1 | 48.179 | 0.425 |
|  | **BON_3** | Ait Ayache |  |  | 14 | 14 | 1 | 0.63 | 0.75 | 0.16 | 0 | 4.78 | 0.16 | 48.179 | 0.425 |
|  | **BON_4** | Ait Bouguemez |  |  | 15 | 15 | 1 | 0.64 | 0.76 | 0.16 | 0 | 4.91 | 0.1 | 48.179 | 0.425 |
|  | **GOT_1** | Gotheron |  | Eco-Ariane & Eco-Melrose & Bio-Smoothie & Eco-Smoothie | 15 | 15 | 1 | 0.64 | 0.73 | 0.11 | 0 | 4.58 | 0.06 | 44.977 | 4.93 |
|  | **GOT_2** | Naour |  |  | 10 | 10 | 1 | 0.63 | 0.75 | 0.16 | 0 | 4.57 | 0.07 | 44.977 | 4.93 |
|  | **GOT_3** | Sighisoara |  |  | 12 | 12 | 1 | 0.65 | 0.74 | 0.12 | 0 | 4.78 | 0.09 | 44.977 | 4.93 |
|  | **GOT_4** | Tauti |  |  | 15 | 15 | 1 | 0.64 | 0.72 | 0.12 | 0 | 4.66 | 0.1 | 44.977 | 4.93 |
|  | **LOO** | Loos-en-Gohelle |  | Jonagored | 15 | 14 | 0.93 | 0.64 | 0.73 | 0.12 | 0 | 4.84 | 0.15 | 50.458 | 2.793 |
|  | **NOI** | Noirlieu |  | Boskoop & Jonagold & Reinette | 13 | 11 | 0.85 | 0.69 | 0.73 | 0.05 | 0 | 4.79 | 0.11 | 48.95 | 4.81 |
|  | **LAB** | Avignon (maintained at ANSES) |  | NA | 10 | 9 | 0.9 | 0.66 | 0.7 | 0.05 | 0 | 4.52 | 0.13 | - | - |
|  | **POI** | Poisy |  | Idared | 7 | 7 | 1 | 0.65 | 0.72 | 0.09 | 0.01 | 4.43 | 0.06 | 45.921 | 6.064 |
|  | **TOU** | Toulenne |  | Val & Akane & Choupette & Topaze | 13 | 13 | 1 | 0.63 | 0.74 | 0.14 | 0 | 4.65 | 0.08 | 44.559 | -0.262 |
| **UK** | **COB** | Cobham |  | 17 different varieties | 16 | 16 | 1 | 0.69 | 0.77 | 0.1 | 0 | 4.81 | 0.08 | 51.399 | 0.392 |

**Table S8 continue.** Genetic diversity statistics, geographic origin, and host species of collected *Dysaphis plantaginea* colonies

| **Country** | **Site** | **Site** | **Host** | **Apple variety** | ***N*** | ***G*** | ***G/N*** | ***H_o_*** | ***H_e_*** | ***F_IS_*** | ***P-value*** | ***Ar*** | ***Ap*** | **Y** | **X** |
| --- | --- | --- | --- | --- | --- | --- | --- | --- | --- | --- | --- | --- | --- | --- | --- |
| **Italy** | **CAD** | Cadriano | *Malus domestica* | Reinette de Champagne | 13 | 13 | 1 | 0.67 | 0.75 | 0.12 | 0 | 4.75 | 0.08 | 44.51 | 11.41 |
|  | **FER** | Ferrara |  | Pink Lady | 12 | 12 | 1 | 0.66 | 0.75 | 0.12 | 0 | 4.92 | 0.38 | 44.87 | 11.49 |
| **Morocco** | **ASN** | Asni |  | Gala Brookfield & Golden & Buckeye | 13 | 13 | 1 | 0.51 | 0.55 | 0.08 | 0 | 3.47 | 0.04 | 31.242 | -7.987 |
|  | **AYA** | Ait Ayache |  | Gala & Golden Delicious & Starking Delicious | 15 | 15 | 1 | 0.51 | 0.61 | 0.15 | 0 | 3.73 | 0.03 | 32.667 | -4.935 |
|  | **BOU** | Ait Bouguemez |  | NA | 15 | 15 | 1 | 0.54 | 0.57 | 0.06 | 0 | 3.63 | 0.02 | 31.646 | -6.468 |
|  | **IMO** | Imouzzer Kandar |  | Gala & Golden Reinders & Golden Smoothee | 9 | 9 | 1 | 0.58 | 0.65 | 0.09 | 0 | 3.9 | 0.1 | 33.731 | -5.016 |
|  | **NAO** | Naour |  | NA | 11 | 11 | 1 | 0.56 | 0.62 | 0.09 | 0 | 3.8 | 0.07 | 32.48 | -5.948 |
| **Romania** | **SIG** | Sighisoara |  | NA | 14 | 14 | 1 | 0.66 | 0.75 | 0.12 | 0 | 4.96 | 0.15 | 46.22 | 24.796 |
|  | **TAU** | Tauti |  | Goldspur & Gloster & Aromat de Vara & Mutsu & William's Pride | 10 | 10 | 1 | 0.66 | 0.74 | 0.11 | 0 | 4.77 | 0.11 | 46.712 | 23.506 |
| **USA** | **GEN** | Geneva |  | Red Delicious & Cortland & Empire & Jonagold | 15 | 14 | 0.93 | 0.57 | 0.64 | 0.11 | 0 | 3.74 | 0.08 | 42.866 | -77.025 |
|  | **WIN** | Winchester |  | Smoothie Golden | 15 | 15 | 1 | 0.58 | 0.63 | 0.09 | 0 | 3.7 | 0.04 | 39.112 | -78.287 |
| **France** | **LOO_P** | Loos-en-Gohelle | *Plantago lanceolata* | NA | 6 | 5 | 0.83 | 0.73 | 0.74 | 0.02 | 0.27 | 4.67 | 0.17 | 50.458 | 2.793 |
| **Spain** | **ALT** | Alta Ribagorça | *Malus sylvestris* | NA | 7 | 7 | 1 | 0.65 | 0.71 | 0.09 | 0 | 4.5 | 0.09 | 42.448 | 0.707 |
| **TOTAL** |  |  |  |  | 667 | 582 | 0.87 | 0.63 | 0.74 | 0.15 | 0 | 4.44 | 0.10 |  |  |

Site: orchard of aphid population sampling; Country: country of aphid population origin; *N*: number of individuals; *G*: number of multilocus genotypes (*MLG*); *G/N*: proportion of unique MLG; *H_O_:* observed heterozygosity; *H_E_*: expected heterozygosity; *F_IS_*: Fixation index; *Ar:* allelic richness; *Ap*: private allelic richness; Y and X: latitude and longitude of the sampling site.

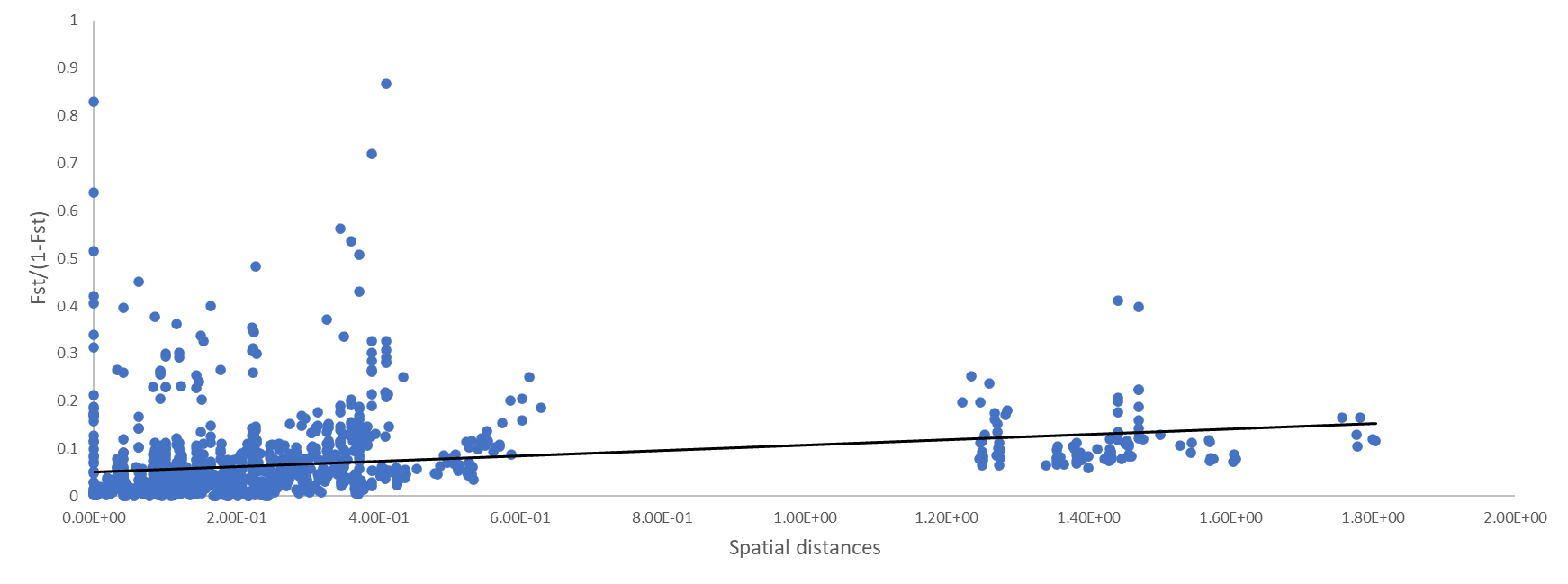

**Figure S11.** Pairwise genetic differentiation (*F_ST_/(1- F_ST)_*) against pairwise spatial distance between the 52 sampling sites for *Dysaphis plantaginea*. All *P-values* were significant.

**Table S9.** Adjusted p-values from the Wilcoxon signed rank test to compare allelic richness (*A_R_*, lower triangle) and private allelic diversity (*A_P_*, upper triangle) among the five *Dysaphis plantaginea* populations detected with STRUCTURE. All comparisons were significant except values in bold.

|  | Spain | Morocco | Eastern Europe & Italy | The US | Western Europe and Denmark |
| --- | --- | --- | --- | --- | --- |
| Spain | 0.000 | **0.47** | **0.38** | 0.016 | 0.00079 |
| Morocco | 9.22e-05 | 0.000 | 0.0042 | **1** | 5.59e-08 |
| Eastern Europe & Italy | 0.196 | 1.99e-05 | 0.000 | 0.000345 | 0.00016 |
| The US | 2.61e-07 | **0.4049** | 2.61e-07 | 0.000 | 5.59e-08 |
| Western Europe and Denmark | 0.00024 | 1.30e-07 | 0.028 | 1.86e-08 | 0.000 |

**Table S10.** Wilcoxon sign-rank test for heterozygosity excess computed using the BOTTLENECK software (Cornuet & Luikart, 1996; Piry, Luikart, & Cornuet, 1999) during range expansion for populations including individuals with a membership coefficient of at least 62.5% to a genetic cluster detected with STRUCTURE for *K*=5.

| **STRUCTURE population** | ***N*** | ***k*** | ***H_e_*** | ***H_eq_*** | ***W1 TPM*** | ***W1 SMM*** |
| --- | --- | --- | --- | --- | --- | --- |
| Western Europe and Denmark | 171.5 | 13.9 | 0.8 | 0.7 | 1.000 | 1.000 |
| Spain | 219.5 | 12.1 | 0.7 | 0.7 | 0.999 | 1.000 |
| Eastern Europe and Italy | 216.9 | 13.3 | 0.7 | 0.7 | 0.998 | 1.000 |
| Morocco | 119.5 | 7.9 | 0.6 | 0.6 | 0.994 | 1.000 |
| US | 51.1 | 5.3 | 0.6 | 0.6 | 0.220 | 0.938 |

TPM = two phase model; SMM = stepwise mutation model; *N* = sample size; *k* = number of alleles; *H_E_* = expected heterozygosity from the observed alleles; *H_eq_* = expected equilibrium heterozygosity; *W1* = Wilcoxon’s signed rank test, probability one tail for *H* excess; TPM: two-phase model; SMM: stepwise mutational model.

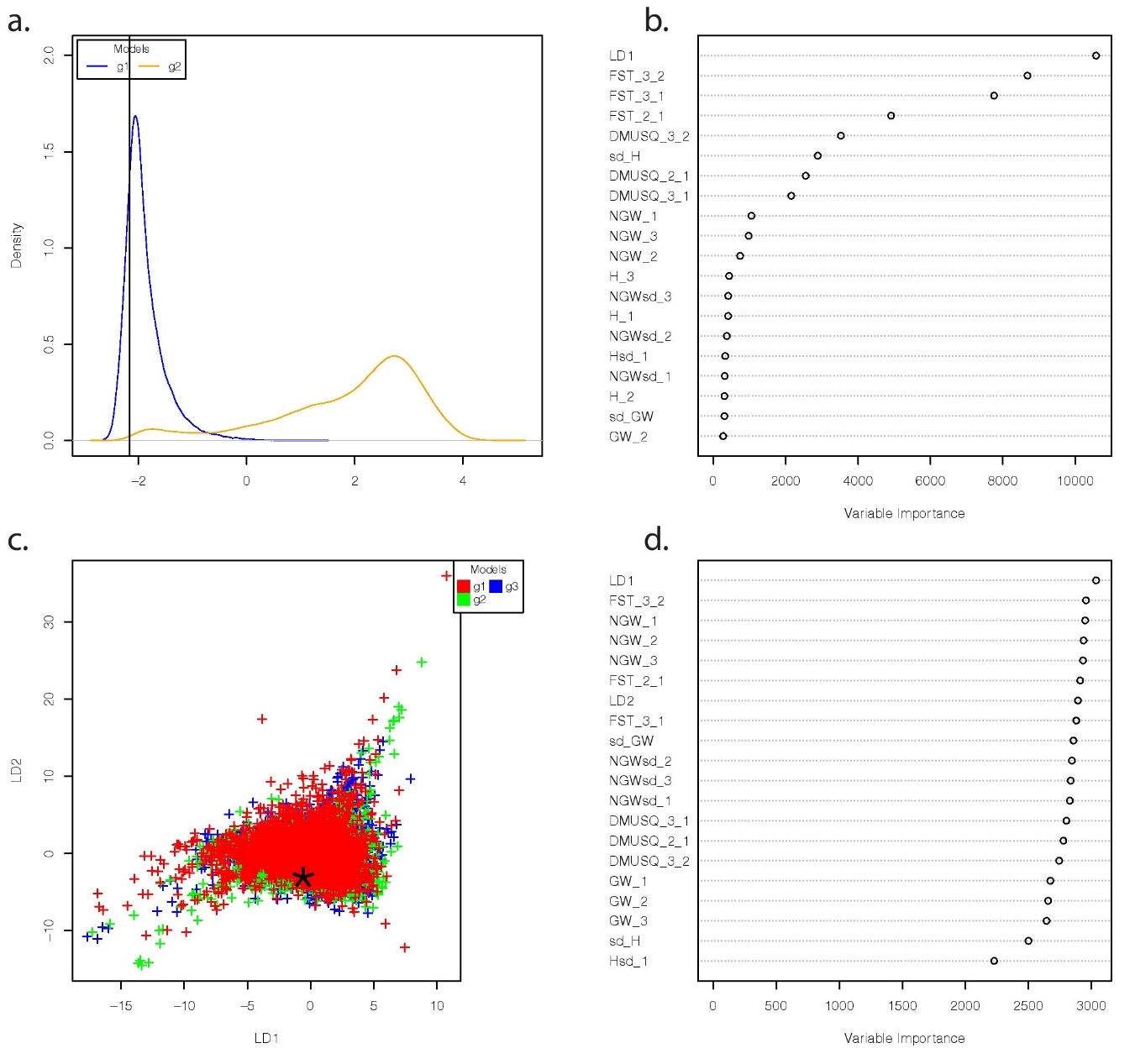
 **Figure S12. Visual assessment of Random-Forest approximate Bayesian computation analyses comparing sequences of colonization of the rosy apple aphid in Europe (ABC-RF step 1) using 29 SSR markers. a.** Projection of the simulated summary statistics obtained from the reference table on a single Linear Discriminant analysis axes for round 1. For round 1, group 1 includes the 24 scenarios depicted in Figure S2a, assuming gene flow among populations, and group 2 includes the same 12 scenarios but assuming no gene flow among populations. Each scenario was simulated 10,000 times. The location of the observed dataset is indicated by a vertical line. The contribution of each statistic is provided in **b.** **c.** Projection of the simulated summary statistics obtained from the reference table on a single Linear Discriminant analysis axes for round 2. For round 2, the groups only include the twelve scenarios assuming gene flow depicted in Figure S2a, and a colonization by the Spanish population (group 1), the Western European/Danish population (group 2), and the Eastern European/Italian population (group 3). The contribution of each statistic is provided in **d.** Each scenario was simulated 10,000 times.

**Table S12.** **Results of the ABC-RF algorithm comparing the colonization history of *Dysaphis plantaginea* in Europe for round 1 (ABC-RF step 1).** Scenarios assumed gene flow among populations (GF) or no gene flow (noGF). We reported in this table: repartition of votes among the two groups of scenarios for each replicate, and mean and standard deviations over replicates for each group of scenarios, posterior probability and prior error rate for the best scenario, *i.e.*, the scenario with the highest number of votes, respectively. The most likely model is highlighted in bold (10 out of 10 votes for GF scenarios).

| Replicate | **GF (group 1)** | noGF (group 2) | Posterior probability | Prior error rate (%) |
| --- | --- | --- | --- | --- |
| 1 | **294** | 206 | 0.64 | 3.0533 |
| 2 | **288** | 212 | 0.62 | 3.0354 |
| 3 | **306** | 194 | 0.60 | 3.0479 |
| 4 | **304** | 196 | 0.56 | 3.0492 |
| 5 | **301** | 199 | 0.64 | 3.0504 |
| 6 | **307** | 193 | 0.62 | 3.0475 |
| 7 | **292** | 208 | 0.59 | 3.0412 |
| 8 | **282** | 218 | 0.61 | 3.0546 |
| 9 | **287** | 213 | 0.61 | 3.0517 |
| 10 | **289** | 211 | 0.57 | 3.0479 |
| Mean | **295** | 205 | 0.61 | 3.04791 |
| sd | 8.88 | 8.83 | 0.023 | 0.006 |

Sd: standard deviation

**Table S13.** **Results of the ABC-RF algorithm comparing the sequences of colonization of *Dysaphis plantaginea* populations in Europe (ABC-RF step 1, round 2).** Scenarios assumed gene flow among populations, as assessed in ABC-RF step 1 - round 1, and a colonization by the Spanish population (group 1), or by the Western European/Danish population (group 2), or by the Eastern European/Italian population (group 3)**.** We reported in this table: repartition of votes among the three groups of scenarios for each replicate, and mean and standard deviations over replicates for each group of scenarios, posterior probability and prior error rate for the best scenario, *i.e.*, the scenario with the highest number of votes, respectively. The most likely model is highlighted in bold (seven out of 10 votes for the group 2, two votes were for the group 1 and one for the group 3). However, note that because of the high prior error rates, those results should be taken with an extra-caution.

| Replicate | group 1  (Spanish) | **group 2 (Western European)** | group 3 (Eastern European) | Posterior probabilty | Prior error rate (%) |
| --- | --- | --- | --- | --- | --- |
| 1 | 146 | **179** | 175 | 0.52 | 60.50 |
| 2 | 151 | **182** | 167 | 0.56 | 60.73 |
| 3 | 168 | **172** | 160 | 0.55 | 60.75 |
| 4 | 188 | **174** | 138 | 0.55 | 60.81 |
| 5 | 157 | **191** | 152 | 0.51 | 60.61 |
| 6 | 152 | **184** | 164 | 0.54 | 60.44 |
| 7 | 162 | **179** | 159 | 0.54 | 60.46 |
| 8 | 165 | **167** | 168 | 0.60 | 60.64 |
| 9 | 185 | **148** | 167 | 0.49 | 60.68 |
| 10 | 176 | **176** | 148 | 0.54 | 60.72 |
| mean | 165 | **175.2** | 159.8 | 0.54 | 60.63 |
| sd | 14.4 | **11.6** | 11.03 | 0.03 | 0.13 |

Sd: standard deviation

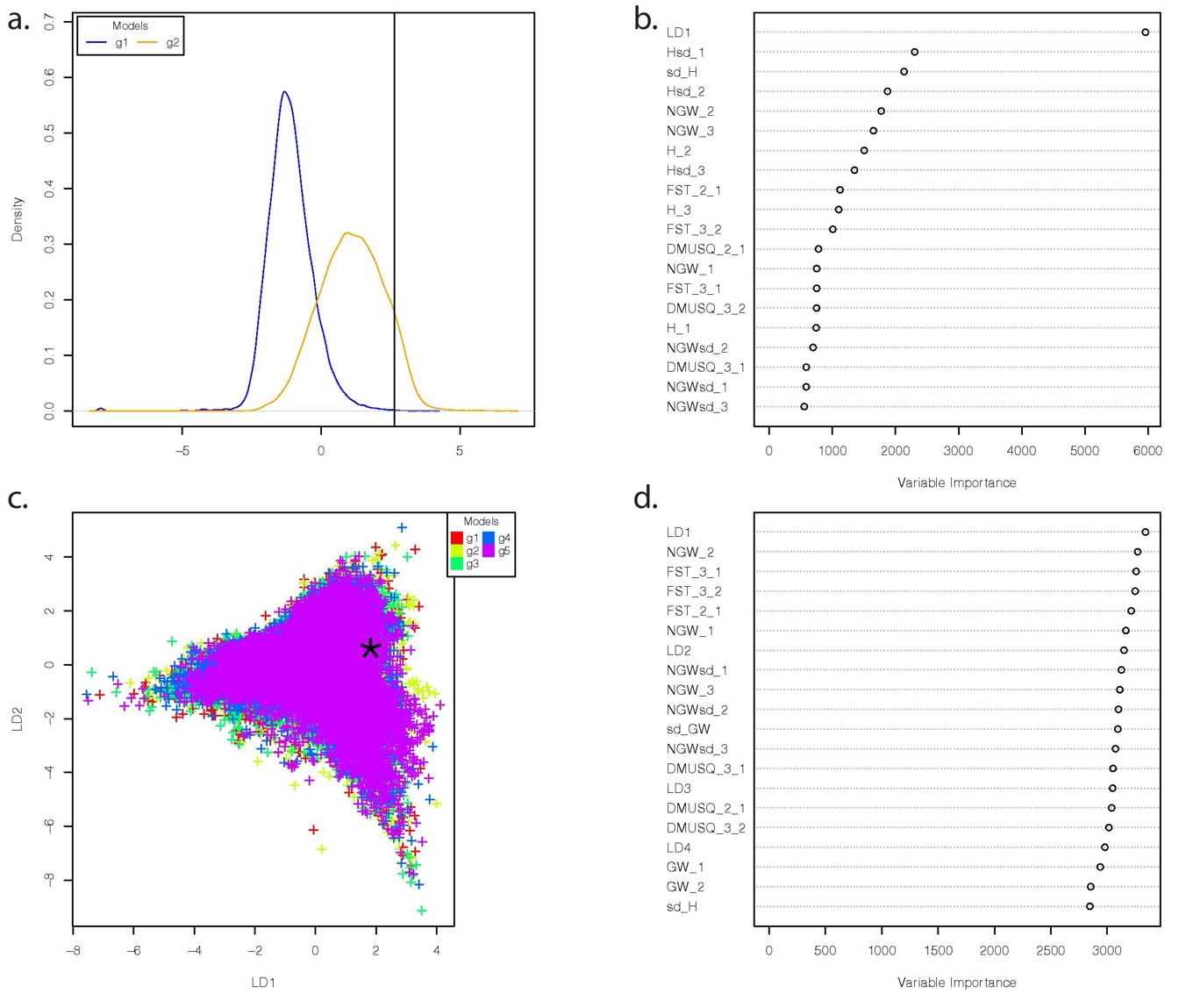
**Figure S13. Visual assessment of Random-Forest approximate Bayesian computation analyses comparing the history colonization of *Dysaphis plantagine*a out of Europe using 29 SSR markers (ABC-RF step 2). a.** Projection of the simulated summary statistics obtained from the reference table on a single Linear Discriminant analysis axes for round 1. For round 1, group 1 includes the six scenarios depicted in Figure S2b, assuming no gene flow among populations, and group 2 includes the same six scenarios but assuming gene flow among populations (*i.e.*, including the four gene flow modalities: all, Moroccan-US, Moroccan-European, US-European). Each scenario was simulated 10,000 times. The location of the observed dataset is indicated by a vertical line. The contribution of each statistic is provided in **b.** **c.** Projection of the simulated summary statistics obtained from the reference table on a single Linear Discriminant analysis axes for round 2. For round 2, each group includes each of the six sequences of colonization depicted in Figure S2b (scenario 13 to 18, respectively); the four gene flow modalities were grouped for a given sequence of colonization (*i.e.*, group). The contribution of each statistic is provided in **d.** Each scenario was simulated 10,000 times.

**Table S14.** **Results of the ABC-RF algorithm comparing the sequence of colonizations of *Dysaphis plantagine*a out of Europe using 29 SSR markers for ABC-RF step 2 (round 1).** Scenarios assumed gene flow among populations (*i.e.*, all, Moroccan-US, Moroccan-European, US-European modalities) or no gene flow**.** We reported in this table: repartition of votes among the three groups of scenarios for each replicate, and mean and standard deviations over replicates for each group of scenarios, posterior probability and prior error rate for the best scenario, *i.e.,* the scenario with the highest number of votes, respectively. The most likely model is highlighted in bold (10 out of 10 votes for the group of scenarios assuming gene flow).

| Replicate | No gene flow (group 1) | **Gene flow (group 2)** | Posterior probability | Prior error rate (%) |
| --- | --- | --- | --- | --- |
| 1 | 151 | **349** | 0.65 | 6.54 |
| 2 | 175 | **325** | 0.66 | 6.55 |
| 3 | 165 | **335** | 0.64 | 6.54 |
| 4 | 155 | **345** | 0.67 | 6.55 |
| 5 | 154 | **346** | 0.66 | 6.55 |
| 6 | 162 | **338** | 0.66 | 6.54 |
| 7 | 172 | **328** | 0.65 | 6.54 |
| 8 | 177 | **323** | 0.65 | 6.56 |
| 9 | 153 | **347** | 0.66 | 6.55 |
| 10 | 159 | **341** | 0.63 | 6.54 |
| Mean | 162.3 | **337.7** | 0.65 | 6.55 |
| Sd | 9.6 | **9.6** | 0.01 | 0.01 |

Sd: standard deviation

**Table S15.** **Results of the ABC-RF algorithm comparing the sequence of colonization of *Dysaphis plantagine*a out of Europe using 29 SSR markers (ABC-RF step 2) for round 2.** Scenarios assumed gene flow (*i.e.*, among all populations, among the Moroccan-US, the Moroccan-European, or the US-European populations), and six different sequences of colonization of the rosy apple aphid (Figure S2)**.** We reported in this table: repartition of votes among the three groups of scenarios for each replicate, and mean and standard deviations over replicates for each group of scenarios; Posterior probability and prior error rate for the best scenario, *i.e.,* the scenario with the highest number of votes, respectively. The most likely models are highlighted in bold (four out of 10 votes for the group 4, four out of 10 votes for the group 2, four out of 10 votes for the group 6). However, note that because of the high prior error rate, those results should be taken with an extra-caution.

| Replicate | g1_scA | g2_scB | g3_scC | **g4_scD** | g5_scE | **g6_scF** | Posterior probability | Prior error rate |
| --- | --- | --- | --- | --- | --- | --- | --- | --- |
| 1 | 78 | 88 | 82 | **92** | 70 | **90** | 0.56 | 73.77 |
| 2 | 69 | 97 | 62 | **96** | 84 | **92** | 0.56 | 73.80 |
| 3 | 84 | 80 | 80 | **98** | 82 | **76** | 0.58 | 73.88 |
| 4 | 70 | 78 | 74 | **91** | 87 | **100** | 0.53 | 73.88 |
| 5 | 90 | 85 | 61 | **87** | 76 | **101** | 0.55 | 73.88 |
| 6 | 75 | 92 | 53 | **109** | 78 | **93** | 0.58 | 73.90 |
| 7 | 79 | 85 | 80 | **95** | 80 | **81** | 0.57 | 73.76 |
| 8 | 88 | 77 | 50 | **107** | 71 | **107** | 0.57 | 73.84 |
| 9 | 87 | 99 | 63 | **77** | 81 | **93** | 0.53 | 73.75 |
| 10 | 76 | 87 | 73 | **97** | 69 | **98** | 0.57 | 73.79 |
| mean | 79.6 | 86.8 | 67.8 | **94.9** | 77.8 | **93.1** | 0.56 | 73.82 |
| sd | 7.4 | 7.5 | 11.6 | **9.2** | 6.2 | **9.3** | 0.02 | 0.06 |

Sd: standard deviation

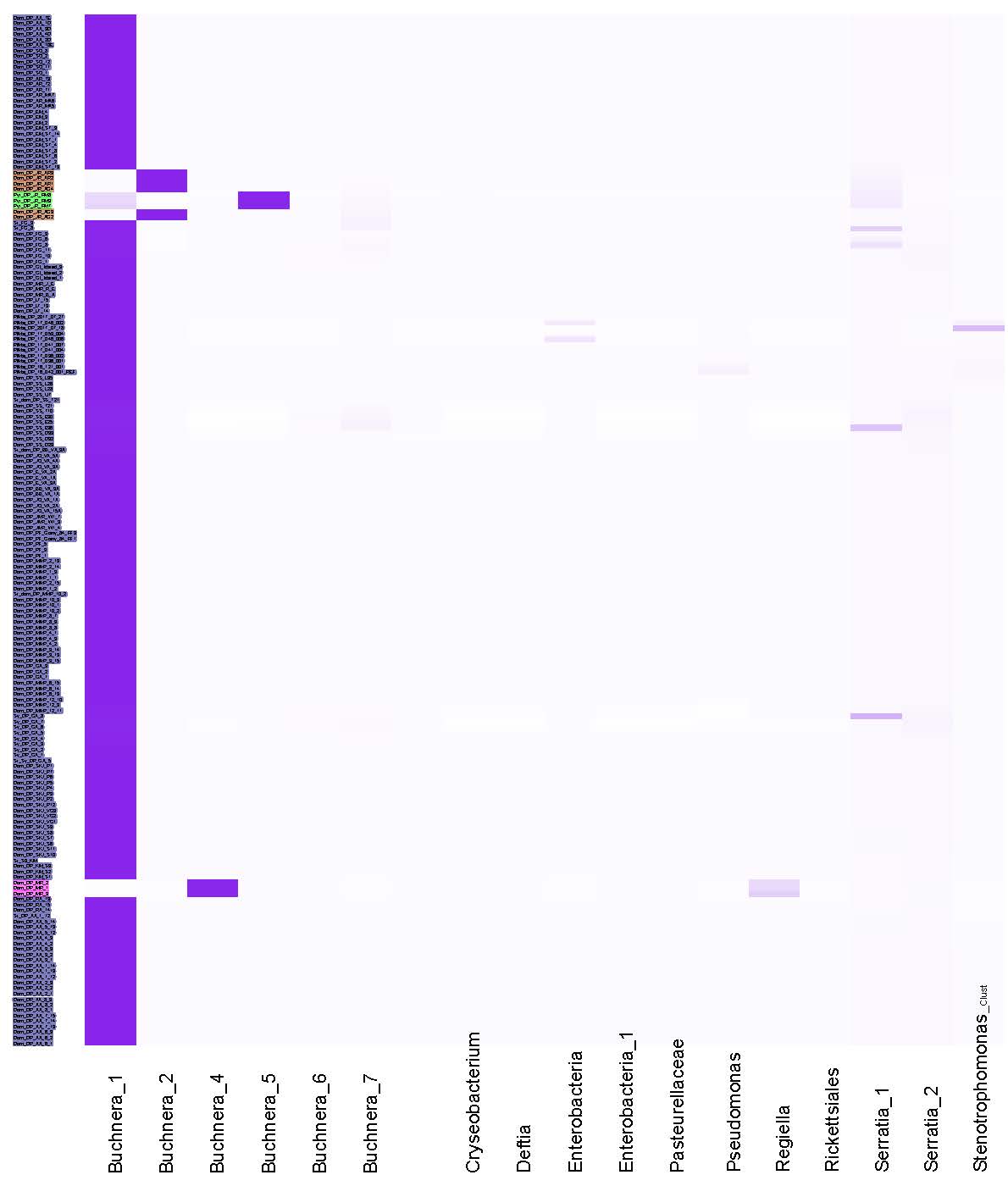

**Figure S14. Low bacterial diversity identified on *Dysaphis plantaginea* across Europe, Morocco and the US.** *Buchnera aphidicola* was identified in most of the *D. plantaginea* individuals (highlighted in blue color). Different *B. aphidicola* groups were detected on *D. plantaginea* from Iran (highlighted in orange color) and *Dysaphis* sp. collected on *Pyrus communis* (highlighted in green color). In addition, other *B. aphidicola* and *Regiella* sp. were observed on *Melanaphis pyraria* (highlighted in pink color). Heatmap of the bacterial diversity of *D. plantaginea* was computed from 5.7 M sequences clustered into 18 OTUs. Each column represents a microbial OTU and color shading indicates the relative abundance. Rows represent each *D. plantaginea* individual. Class level designation for the microbial OTUs is presented at the bottom of the Figure.
