## Supporting_information text S1 for "Large-scale geography survey provides insights into the colonization history of a major aphid pest on its cultivated apple host in Europe, North America and North Africa"

beyond

### **Text S1. Protocol for DNA extraction of *Dysaphis plantaginea***

**Each colony was kept in ethanol (96%) until DNA extraction:** One individual per colony to perform DNA extraction. The night before the DNA extraction one individual was picked up from the colony kept in 96% ethanol and put at -80°C. The next day the aphid was grounded to fine dust by two cycles of cryo-grinding of the 30s at 23Hz.

**Extraction step:** 300µL of extraction buffer containing TNES buffer (50 mM Tris pH 7.5, 400 mM NaCl, 20 mM EDTA, 0.5% SDS, water), proteinase (50 µg) and RNAse (100µg) was then added to the tube containing aphid dust, before being incubated at 55°C for 1 hour with 1000 rpm agitation.

**Deproteinization and filtration steps**: We then added 85µL of NaCl (5M) to the mix, shaking vigorously for 15 seconds. Tubes were then centrifuged 20 minutes at 4°C and 5000 rpm.

**DNA fixation steps:** 300 µL of supernatant were then transferred to a Whatman plate mounted on a deep well plate, to which we added 500 µL of CGE buffer (2.6 M guanidinium chloride, water, 64% ethanol). The plate was then centrifuged for 2 minutes at 5000 rpm at room temperature. The eluate was discarded.

**Washing step:** We then proceeded to two consecutive washes. A first wash with 600 µL of washing buffer (8mM Tris pH 8, 0.04 mM EDTA, 60 mM potassium acetate, and 60% ethanol) and with two minutes of centrifugation at 5,000 rpm at room temperature; eluate was discarded. The second wash consisted of the same amount of washing buffer with 15 minutes of centrifugation at 5,600 rpm at room temperature. Eluate was discarded.

**DNA elution step:** The Whatman plate was then put on top of a PCR plate. A total of 30 µL of 65°C water was added to each column. After 5 minutes the plate was then centrifuged for 2 minutes at 1000 rpm at room temperature. This step was repeated twice.

### **Text S2. Construction and sequencing of microsatellite enriched libraries**

We developed 29 new SSR markers using a low coverage genome. The *D. plantaginea* low-coverage genome sequenced in this study originated from a pool of genomic DNA of five genotypes from five colonies from three distinct locations in Europe with a homemade routine DNA extraction protocol (Text S1). We used a total of 1 µg from an equimolar DNA pool of the five genotypes for the development of a DNA genomic library. Preparation of DNA libraries was performed by the Genoscreen company (Lille, France [www.genoscreen.fr)](about:blank), which then used the Illumina MiSeq platform to generate 300-bp paired-end reads. A total of 1,157,582 reads were thus obtained.

Genoscreen used the algorithm implemented in the Velvet package (Zerbino and Birney, 2008) designed to deal with *de novo* genome assembly and short-read sequencing alignments. The assembling was achieved by testing k-mers from 31 to 121 with a step of 10. The assembling with the highest rate of remapping was kept. The bioinformatics program QDD v3 (Meglécz et al., 2014) was used to analyze sequences. QDD treats all bioinformatics steps from raw sequences until obtaining PCR primers: removing adapters/vectors, detection of microsatellites, detection of redundancy/possible mobile element association, selection of sequences with target microsatellites and primer design by using BLAST (<https://blast.ncbi.nlm.nih.gov/Blast.cgi)>, ClustalW (Larkin et al., 2007) and Primer3 (Rozen and Skaletsky, 2000) programs. The 1,157,582 reads were therefore assembled in 50,445 contigs.

From the 50,445 obtained contigs with QDD, sets of microsatellite primers were designed on suitable microsatellite ﬂanking regions. From this set of microsatellite primers, the Genoscreen company only kept perfect di/tri/tetra motifs, with A and B designs (according to QDD internal parameters: regions that do not have multiple microsatellites, nanosatellite, and homopolymers) and a minimum of 20 bp between the primer and the microsatellite. A total of 4,380 primer sets were designed on suitable microsatellite ﬂanking regions, and 126 primers pairs were initially selected as being the more promising by the Genoscreen company. Out of those 126 primers, a total of 33 primers, were then tested for amplification and polymorphism tests.

### **Text S3. Polymorphism screening of the microsatellite markers and panel construction**

#### **Material and methods**

As a matter of comparison with a previous study that has investigated *D. plantaginea* population structure (Guillemaud et al., 2011), amplification and polymorphism were also tested on seven previously developed SSR markers.

We, therefore, tested 40 SSR for polymorphism screening, including 33 newly developed and seven previously developed SSR markers (Guillemaud et al., 2011). For screening microsatellite polymorphism for *D. plantaginea*, we selected eight individuals from distinct geographical locations in Europe: France (two samples), Spain (two samples), Denmark (one sample), Germany (one sample), Czech Republic (one sample), Belgium (one sample). DNA was extracted following the protocol described in Text S1. PCR amplifications were performed for each microsatellite marker with unlabeled specific primers using a home-made Taq DNA Polymerase and temperatures of hybridization calculated based on the primer sequence. Amplification success was then tested on electrophoresis (size control) and SSR polymorphism screened on acrylamide gels. We only retained SSR markers based on their range of allelic size, and polymorphism. Retained SSR markers showed 1) PCR products that migrated into the electrophoresis gel for a size corresponding to the size predicted with the Primer3 software, 2) at least two alleles scored on the eight reference individuals on the acrylamide gels, 3) clear bands on the gel, *i.e.*, without non-specific bands.

We tested the neutrality for microsatellite markers with the Ewens-Watterson neutrality test (Watterson, 1978); the statistics *F* (sum of the square of allelic frequency) and limit (upper and lower) at 95% confidence region for the test were calculated using the algorithm by using 1,000 simulated samples implemented in the Popgene software (Yeh et al., 1997). The markers could, therefore, be considered unlinked and neutrally evolving.

Each panel of microsatellite markers was then constructed including four markers of non-overlapping allelic sizes to avoid allele size overlapping during genotyping. Each panel contained four markers tagged with four different fluorescent dyes (ATTO 550, ATTO 565, HEX, and Alexa Fluor, Table S4 in the main supplementary file).

#### **Results**

Four out of the seven previously developed SSR markers did not pass the first amplification validation step. The Sa_4s, Sav_S3.43, Sm_S16b, and Rp_R5.29B markers indeed showed either low polymorphism or non-specific bands on electrophoresis gels (data not shown). We, therefore, discarded those four SSR markers.

We therefore kept 36 SSR for polymorphism screening with acrylamide gels, including 33 newly developed and three previously published SSR markers. The screening of the genotypes was difficult to assess with certainty for six markers out of the 36 initial SSR. Those six SSR markers were therefore discarded from the dataset, leading us to keep 30 SSR markers including 29 newly developed SSR and one previously published marker (DpL4) (Guillemaud et al., 2011). Details on primer sequences, motifs, and repeat number, range, panel, dye, the temperature of hybridization, alleles, observed and expected heterozygosity, and fixation index for each of the 30 microsatellite markers can be found in Table S5 (main supplementary file). We further checked the suitability of the 30 microsatellite markers for population genetic analyses. None of the 30 microsatellite markers deviated significantly from a neutral equilibrium model (Table S4, main supplementary file); none F-value (sum of the square of allelic frequency) lied outside the lower and upper limit of 95% confidence region of expected F value for any loci, and no pair of markers was found to be insignificant linkage disequilibrium in any of the species.

We further tested the polymorphism of the 30 retained microsatellite markers retained. The number of alleles per locus varied from 8 to 50, for a total of 640 alleles observed. Descriptive statistics on the polymorphism of *D. plantaginea* and on deviations from Hardy-Weinberg equilibrium can be found in Table S5 (main supplementary file). The mean observed heterozygosity (*H_O_*) was 0.63 with values ranging from 0.32 to 0.91, and the mean expected heterozygosity (*H_E_*) was 0.74 with values ranging from 0.46 to 0.92. The fixation index (*F_IS_*) ranged from -0.006 to 0.65 with a mean value of 0.15.
